# Protein architectures of the bacterial spore envelope - common principles of assembly

**DOI:** 10.64898/2026.09.03.749163

**Authors:** Hannah Fisher, Abigail R. E. Roberts, Ainhoa Dafis-Sagarmendi, Nic Mullin, Jessica E. Buddle, Thamarai K. Janganan, Jamie K. Hobbs, Robert P. Fagan, Per A. Bullough

## Abstract

Bacterial spores are among the most durable of cell forms, protected by robust envelopes of layered protein assemblies. Remarkably, these envelopes often share similar architectures across distantly related bacteria despite extensive divergence in their molecular components. How evolution has converged on such similar highly ordered and resilient cellular structures is relatively unexplored. Here, we combine targeted mutagenesis, cryo-electron microscopy, atomic force microscopy and structure prediction to determine the organisation and assembly of the outer spore envelope of *Clostridium sporogenes*, a genetically tractable surrogate for Group I *Clostridium botulinum*. We reveal a hierarchy of protein structures, including a semi-permeable two-dimensional crystalline exosporium; CsxA forms the exosporium scaffold with an outer “hairy nap” partially composed of BclA. We observe a previously uncharacterised multilayered three-dimensional crystalline parasporal assembly within the interspace between exosporium and coat (CsxC). These distinct structures are built from related cysteine-rich SPOCS (SpoVID-CotE-SipL)-domain proteins that self-assemble into highly ordered lattices. Such crystalline organisation, combined with high symmetry, can provide a template for local enhancement of cysteine concentration, favouring cooperative disulphide cross-linking and the formation of exceptionally stable supramolecular structures. Unexpectedly, the three-dimensional CsxC structure grows through screw dislocations, a mechanism familiar from inorganic and synthetic crystal growth but rarely demonstrated in native biological assemblies, with the exception of some biomineralisation processes. Our observations now suggest that this classical crystal-growth mechanism can be exploited in both mineralised and proteinaceous biological materials. Related SPOCS-domain proteins are implicated in spore-envelope assembly across diverse Clostridia, where they have diversified to act as both structural components and morphogenetic organisers. Remarkably, distantly related Bacilli construct similarly highly symmetric crystalline, cysteine-rich and disulphide-stabilised spore layers using proteins with very different protein folds. Thus, crystallisation and cooperative disulphide formation appear to represent a convergent physicochemical strategy for building diverse self-assembling proteins into exceptionally robust cellular assemblies. Together these findings provide the first molecular framework for understanding the organisation and assembly of the *Clostridium* spore envelope and reveal previously unrecognised principles governing the evolution of protective proteinaceous structures across the Bacillota (Firmicutes).

## Introduction

Endospore (spore) formation in the Bacillota provides a striking example of both conservation and divergence. The core regulatory mechanisms controlling sporulation are deeply conserved (Galperin et al., 2022), yet the proteins forming the outermost protective layers of spores show remarkable divergence between species. Despite this, apparent convergent evolution in distantly related spore-forming species has led to outer envelopes with often similar morphology and properties: highly ordered, exceptionally resistant layered protein assemblies. How are these similar biological structures constructed from such different molecular components?

Spores are among the most durable of cells (Setlow, 2014b). Their remarkable resilience underpins the environmental persistence and transmission of medically and economically important members of the Bacillota, including *Bacillus anthracis*, *Clostridioides difficile* and proteolytic Group I *Clostridium botulinum* (Koch 1876; Arnon, 1977; Sorg and Sonenshein, 2008). The envelope of the spore, which forms the interface between the dormant cell and its environment, plays central roles in this resilience, along with protection, environmental sensing and the initiation of germination (Driks 2016; Henriques 2007; Setlow, 2014a). It is a hierarchically organised, layered structure comprising the cortex, protein coat and, in many species, an additional outermost layer known as the exosporium (McKenney 2013).

The molecular architecture and function of the exosporium has only been characterised in detail in the *Bacillus anthracis/cereus/thuringiensis* group; it consists of a crystalline basal layer decorated by a collagen-like hairy nap. It provides one of the best characterised examples of a self-assembling cellular protein superstructure (Boone et al., 2018; Henriques and Jr., 2007; Ball et al., 2008; Kailas et al., 2011; Stewart, 2015; Stewart, 2017; Terry et al., 2017; Janganan et al., 2020; Portinha et al., 2022). Similar organisational principles appear to operate in Clostridia, but the molecular details are still largely unknown (Janganan et al., 2020; Hoeniger and Headley, 1969; Mackey and Morris, 1972; Lund et al., 1978; Masuda et al., 1980).

Spore envelopes of Bacillota are often morphologically similar to each other (Ball et al., 2008; Rabi et al., 2017; Hoeniger and Headley, 1969; Driks and Eichenberger, 2016) yet comparatively few coat or exosporium proteins are shared between them (Galperin et al., 2012; Galperin et al., 2022; Davidson et al., 2018; Secaira-Morocho et al., 2020). This raises a fundamental evolutionary question: have these organisms constructed their similar protective envelopes using structurally homologous assemblies despite sequence diversion, or have they independently evolved distinct structural solutions to the same protective challenge? Recent studies suggest that cysteine-rich proteins may represent a recurring solution for constructing these highly stable spore outer layers (Jiang et al., 2015; Terry et al., 2017; Stewart, 2017; Janganan et al., 2020; Romero-Rodríguez et al., 2020). Despite displaying little sequence similarity, these proteins frequently contain high concentrations of cysteine residues predicted to stabilise supramolecular assemblies through co-operative disulphide bonding. However, studies of the structures of these assemblies have mostly been confined to the Bacilli; for Clostridia, only one case has been studied in detail (Janganan et al., 2020).

Beyond *Bacillus subtilis* and the *Bacillus cereus* group, *Clostridium sporogenes* provides an excellent model system for addressing questions of spore envelope assembly. As a close non-toxigenic relative of proteolytic Group I *C. botulinum*, it produces spores that are morphologically similar while remaining genetically tractable and easily handled in the laboratory (Portinha et al., 2022; Masuda et al., 1980; Fisher et al., 2026). We previously identified three cysteine-rich proteins, CsxA, CsxB and CsxC, together with the collagen-like proteins BclA and BclB, as candidate structural components of the outer spore envelope (Janganan et al., 2016). That work also revealed several distinct classes of filamentous spore appendages but their site of attachment and relationship to the envelope architecture remain unknown. Recombinant CsxA self-assembles into highly ordered two-dimensional crystalline arrays essentially identical to the exosporium basal layer, identifying it as the principal structural component of that layer (Janganan et al., 2020). However, the functions of the remaining proteins, the molecular organisation of the complete envelope and the relationship between these proteins remain unresolved.

Here, we combine genetic dissection with thin-section, cryo-electron, and atomic force microscopy to determine how the outer spore envelope of *C. sporogenes* is organised from the cellular to the molecular scale. We show that the envelope comprises genetically and structurally distinct crystalline assemblies and define the contributions of the cysteine-rich proteins CsxA, CsxB and CsxC to their organisation. We identify a novel mechanism of three-dimensional crystal growth amongst these proteins. Comparison of these Clostridial assemblies with those of other spore-forming bacteria further suggests that similar spore envelope structures have emerged from very different molecular building blocks. These findings provide a framework for understanding both the physico-chemical principles governing assembly of the spore envelope and its evolutionary diversification.

## Results

### Overall architecture of the dormant *Clostridium sporogenes* spore envelope

The exosporium is a loose, bag-like structure that surrounds, and is approximately twice the length of, the main spore body. Although the spore body could potentially occupy different positions within the exosporium, in more than 70% of wild-type spores it is positioned close to one pole (Fig 1A). Internally, the spore body consists of a core, containing the genetic material, surrounded by the cortex and proteinaceous coat (Fig 1B-C). The spore body is separated from the exosporium by the intervening interspace, containing prominent laminated structures that are morphologically distinct from both the coat and exosporium. Thicker stacks of these layers tended to localise around the equatorial region of the spore (Fig 1B); by analogy with *B. cereus* group spores (Wehrli et al., 1980; Ebersold et al., 1981; Ball et al., 2008), we refer to these assemblies as **parasporal layers**.

**Fig 1.**
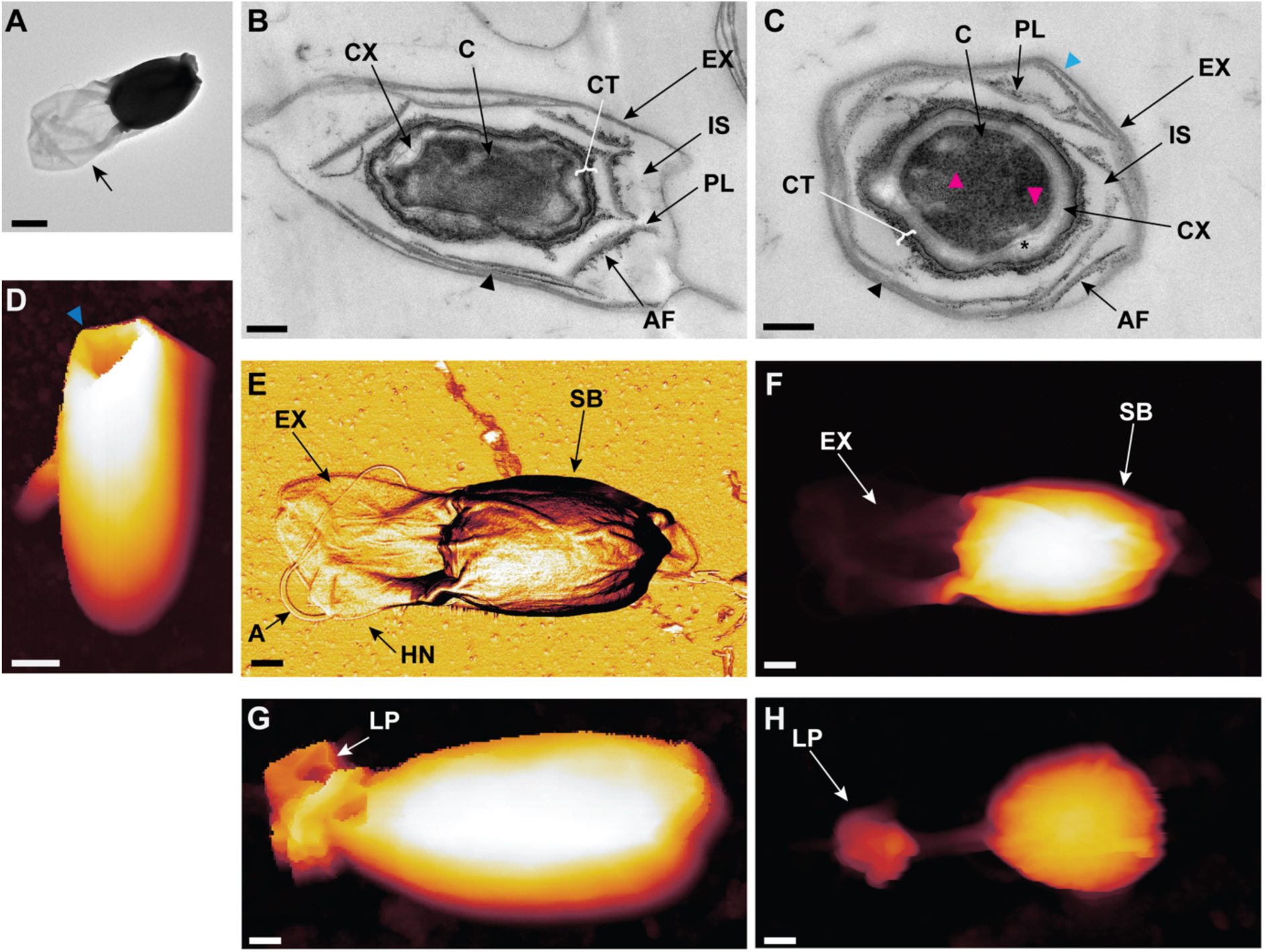
Overall architecture of the dormant *Clostridium sporogenes* spore envelope. **A** Representative electron micrograph of negatively stained spore (Strain NCIMB 701792 wild type) (n = 73). The spore body occupies one pole of the exosporium, leaving the other pole unoccupied. (black arrow). Scale bar: 0.5 µm. **B**-**C** Representative thin sections of WT spores (n = 32), sectioned parallel to the polar axis (B) and perpendicular to the polar axis (C). Labels: C - core, CX - cortex, CT - coat, IS - interspace, PL - parasporal layer, AF - amorphous fringes, EX - exosporium, black arrow heads - parasporal layer laminations, magenta arrow heads - ribosomes, light blue arrow heads - crystalline lattice. Scale bar: 200 nm. **D** Atomic force micrograph (height image) of spore in liquid showing an aperture at one pole of the exosporium - dark blue arrowhead. Height range: 0 - 1.5 μm. Scale bar: 500 nm. **E** Atomic force micrograph (phase image) of dormant spore in air, showing polar arrangement of the exosporium relative to the spore body (SB), hairy nap (HN), and appendages (A). Scale bar: 250 nm. Phase range: 21.9 degrees. **F** Atomic force micrograph (height image) of the dormant spore (E), showing extended polar end of the exosporium. Scale bar: 250 nm. Height range: 0 - 900 nm. **G-H** Atomic force micrograph (height image) of dormant spore in liquid (G) and air (H) showing a lipped terminal protrusion (LP). Scale bar: 250 nm. Height range: 0 - 1 μm.

High-pressure freezing and freeze-substitution, which avoid potential artefacts arising from conventional chemical fixation, confirmed that the parasporal layers are common features of the spore (Fig 2). These tended to stack together forming laminations abutting the interior face of the exosporium basal layer but were also seen as detached fragments within the interspace, decorated on one side with a darkly stained amorphous fringe (Fig 1B-C). Transverse sections were consistent with stacks being wrapped cylindrically around the spore body (Fig 2). The periodicity of the laminations measured from Fig 2 in their stacking direction was ∼55 Å (average over four layers) and the periodicity in the plane of each lamination was measured at ∼50 Å (average over ten repeats). Tomographic reconstruction further demonstrated that the lamellae occupy the interspace between the coat and exosporium as a coherent three-dimensional assembly (Fig 2, S1 Fig, S1 Movie).

**Fig 2.**
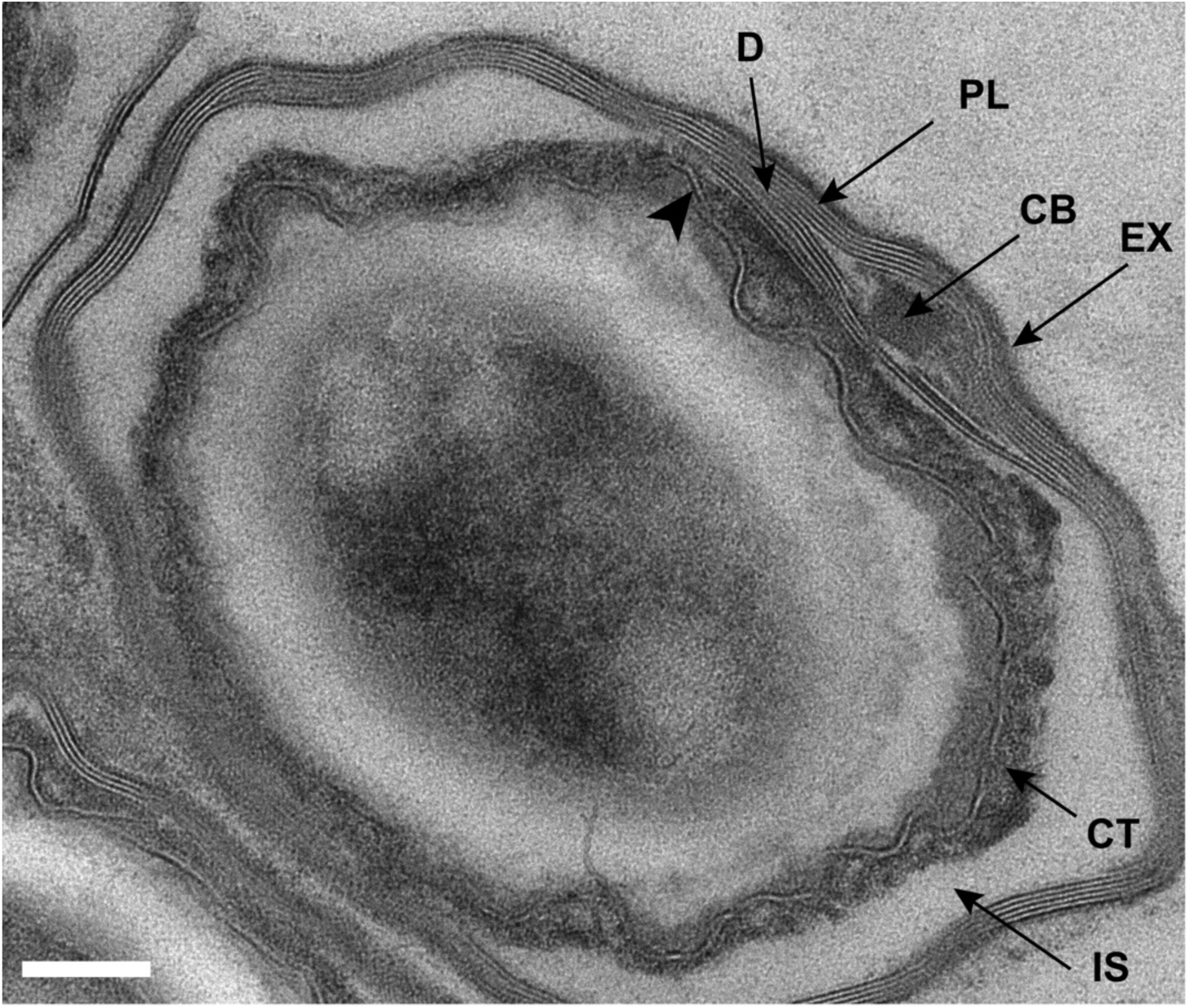
High-pressure freezing and freeze substitution preserves the multilamellar parasporal layers occupying the interspace of the native spore envelope. The hirsute exosporium (EX) forms the outer surface. Apposed to this are multiple lamellae of the parasporal compartment (PL) which display discontinuities (D). In some, but not the majority of sections we saw the layers wrapped around an unidentified cuboid body (CB). The exosporium and parasporal layers are separated from the spore coat (CT) by an intervening interspace (IS). In this sample preservation method the coat is seen to be made up of multiple differentially staining layers. The central pale band displays hints of a regular array in places (arrowhead). This level of detail has not been previously reported for conventional resin embedded samples (Fig. 1) (Walker et al., 1967; Hoeniger and Headley, 1969; Stevenson and Vaughn, 1972; Brunt et al., 2015). Scale bar: 100 nm

AFM imaging in air confirms the polar arrangement of the exosporium relative to the spore body; this collapses onto the substrate surface when away from the spore body during drying. When fully hydrated in liquid the exosporium forms an inflated structure, hiding the spore body from view. A dense hairy nap is revealed to cover the spore surface (Fig 1E-F). At the extended exosporium pole, two morphologies were frequently observed-a third of spores displayed an apparent circular opening, whereas about a fifth displayed a lip-like cap enclosing the polar extension (Fig 1D, G-H).

### Genetic dissection reveals distinct roles for CsxA, CsxB and CsxC in spore envelope assembly

To determine the contributions of the three cysteine-rich proteins to spore envelope architecture, we examined the morphology of Δ*csxA*, Δ*csxB* and Δ*csxC* spores and compared them with wild-type (Fig 3). The three mutations produced distinct phenotypes that genetically separate formation of the exosporium from assembly of the parasporal layers within the interspace.

**Fig 3.**
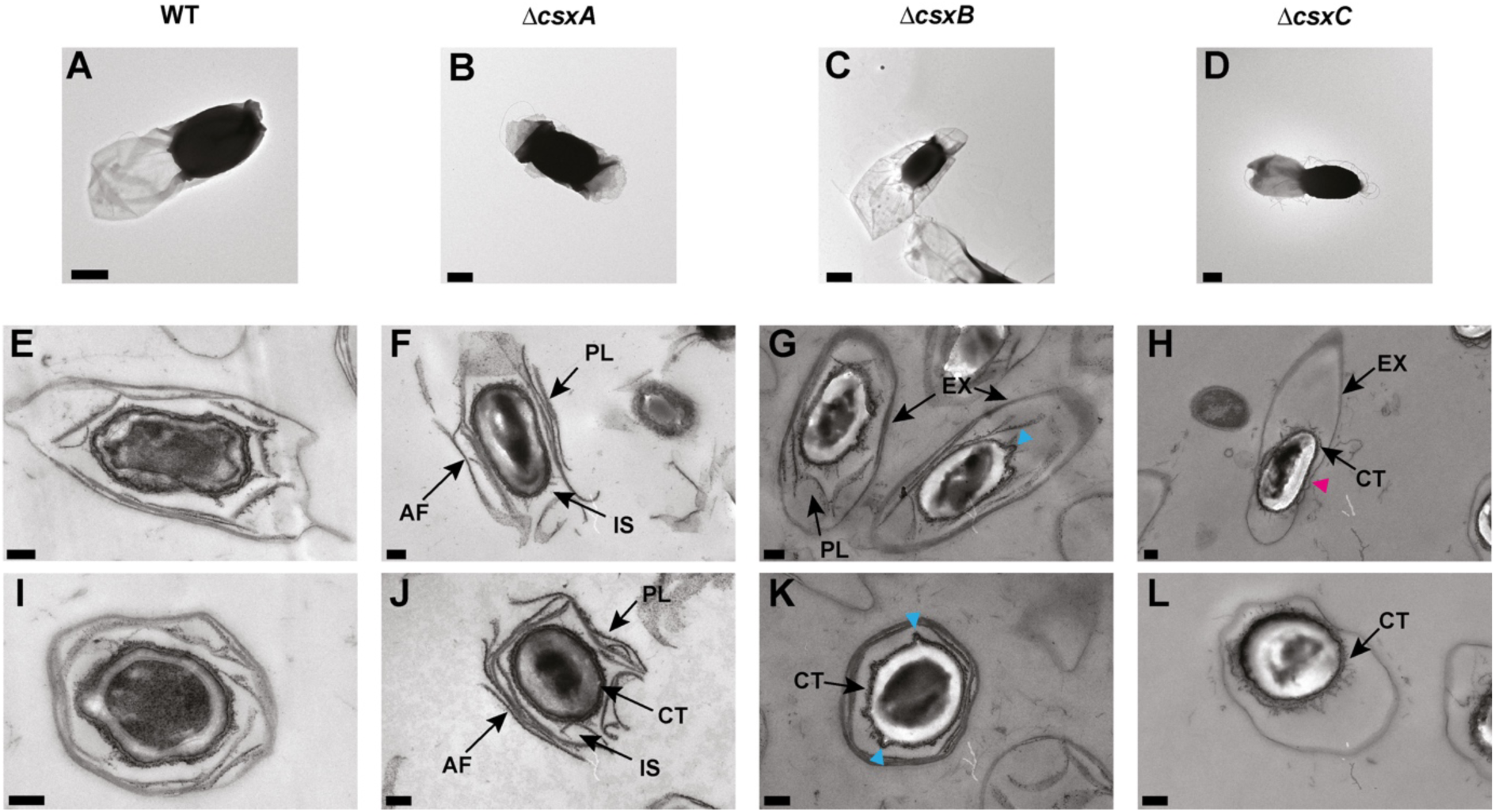
Genetic dissection reveals distinct roles for CsxA, CsxB and CsxC in spore envelope assembly. **A-D** Representative micrographs of negatively stained WT, Δ*csxA*, Δ*csxB* and Δ*csxC* (Strain: NCIMB 701792) spores. Scale bars: 0.5 µm. **E-H** Representative corresponding micrographs of thin sectioned resin embedded spores sectioned approximately parallel to the polar long axis. Scale bars: 200 nm. **I-L** as (E-H) but presumed to be sectioned approximately perpendicular to the polar long axis. Scale bars: 200 nm Features of note are labelled as follows: PL - parasporal layers, IS - interspace, CT - coat, EX - exosporium, light blue arrow heads - coat blebs, magenta arrow head - exosporium tight to the coat. Negative Stain: WT n = 73 spores Δ*csxA* n = 17 spores, Δ*csxB* n = 11 spores, Δ*csxC* n = 8 spores, Thin Sections: WT n = 32 spores Δ*csxA* n = 34 spores, Δ*csxB* n = 29 spores, Δ*csxC* n = 27 spores.

#### CsxA is essential for exosporium assembly

Deletion of *csxA* resulted in the production of spores that completely lacked the characteristic balloon-like exosporium (Fig 3B, 4A). The cortex, coat, interspace and parasporal layers appeared largely intact and stacks of parasporal layers surrounded the spore body (Fig 3F, J). Thus, CsxA is required for exosporium assembly but is not essential for the assembly of the main spore body or for the formation and retention of the parasporal layers.

Consistent with this, recombinant CsxA self-assembles into highly ordered two-dimensional crystals indistinguishable from the native exosporium basal layer, providing an opportunity to analyse the hydrated lattice by cryoEM (Janganan et al., 2020). Combining cryoEM with AlphaFold3 structure prediction allowed us to propose a molecular interpretation of the lattice.

#### The exosporium is built from a continuous hexagonal protein lattice

Three-dimensional electron crystallography of tilted frozen-hydrated CsxA crystals yielded structure factors to 9 Å resolution in plane and 20 Å perpendicular to the plane. Data are summarised in S1 Table. CsxA assembles into a continuous two-dimensional crystalline array composed of hexagonally arranged crown-like assemblies interconnected by trimeric linkers (Fig 4A-B, S2 Movie). This resembles the *in situ* exosporium basal layer arrangement we described previously (Janganan et al., 2020). The crown-like assemblies surround a central pore, while the connecting trimeric linkers generate an interconnected network of contacts throughout the crystal. Peripheral pores are present in between these linkers, ovoid in shape with a width of ∼15 Å and a length of ∼45 Å.

**Fig 4.**
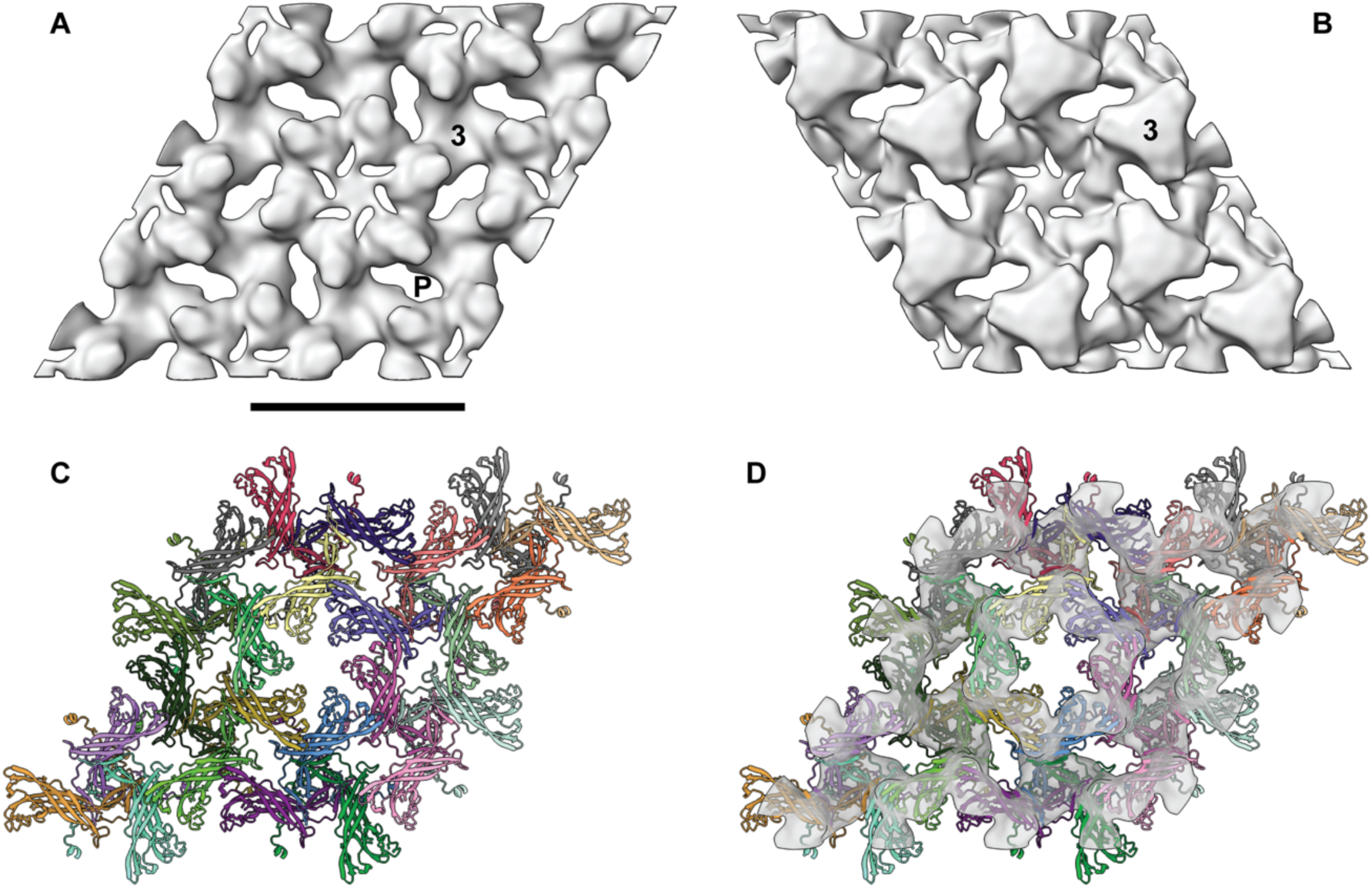
CryoEM reveals the molecular architecture of the CsxA exosporium lattice. **A** Reconstruction with the threshold set approximately to a volume corresponding to 33.4 kDa protein per subunit (assuming protein density of 1.35 g/cm^3^). Noise is enhanced sixfold on the central symmetry axis so this density is unlikely to be significant. 3- threefold linker, P- blate pore. Scale bar: 105 Å. The view is predicted to be onto the external spore face (see Fig. 9). **B** Equivalent view onto the internal face. **C** and **D** Molecular model of the array of individual CsxA subunits coloured (see Fig 7B). The map threshold has been raised to reveal the central pore.

#### Structural modelling of CsxA provides a molecular interpretation of the cryoEM density

To interpret the cryoEM reconstruction, CsxA was modelled using AlphaFold3 (Abramson et al., 2024), with a trimeric arrangement emerging as the most confident prediction. Foldseek (Kempen et al., 2024) suggested that CsxA is a member of the Pfam SPOCS (SpoVID-CotE-SipL) domain superfamily incorporating the Pfam DUF3794 fold (Paysan-Lafosse et al., 2024; Delerue et al., 2022) (S2, S5 Fig). In the predicted trimer the N-terminal proximal sheet of chain A is extended by one additional β-strand from chain B (Fig 5). Each subunit contains 24 cysteine residues, several of which are positioned in the AlphaFold3 model such that disulphide bonds could feasibly form. (S2, S5 Fig, S1 Table). Such cross-linking could contribute to stabilisation of the crystalline assembly (Janganan et al., 2020).

**Fig 5.**
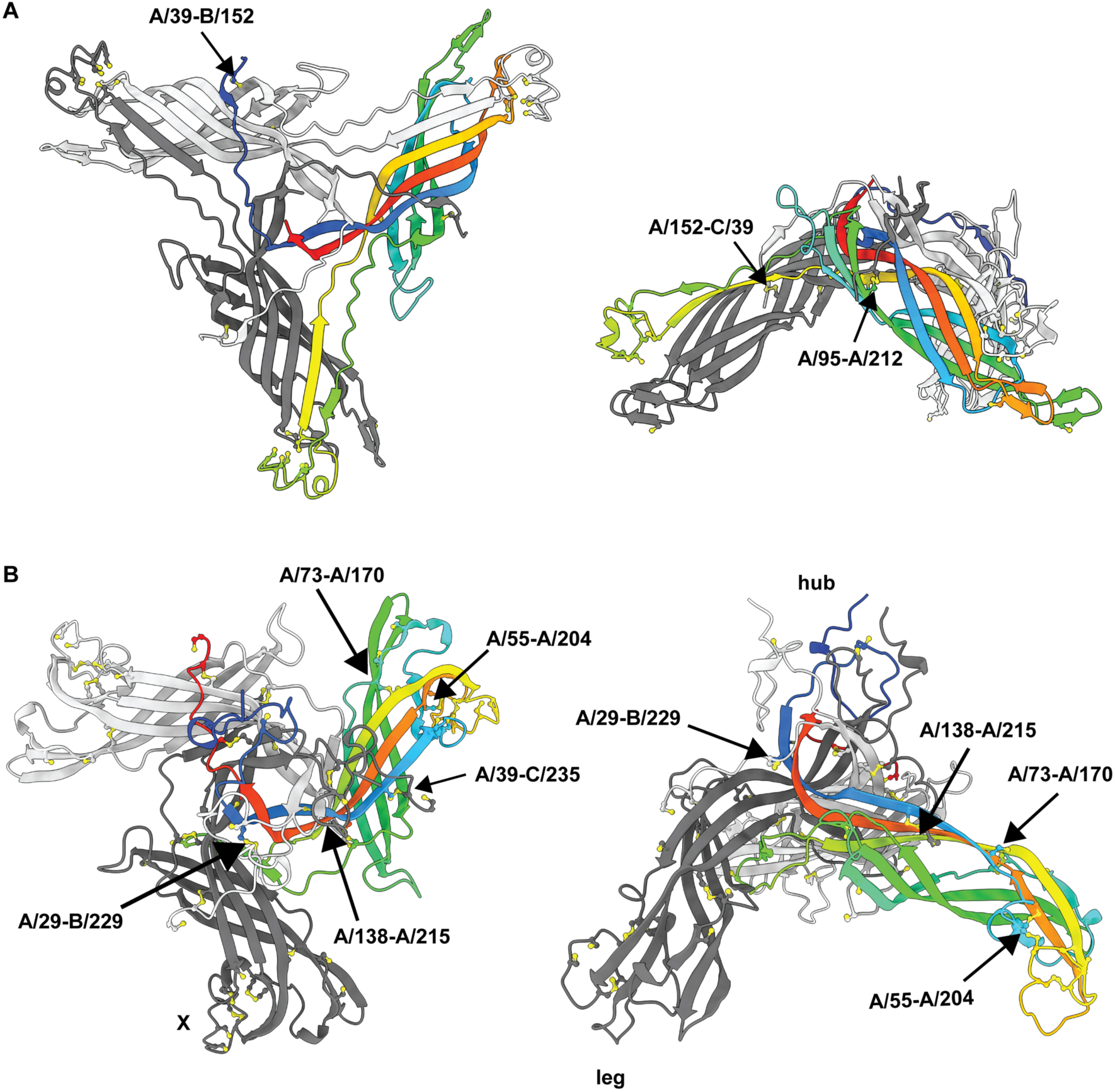
AlphaFold3 predicts that CsxA and CsxC share a conserved cysteine-rich trimeric scaffold. **A** Two views of CsxC with one subunit (A) coloured from N-terminus (blue) to C-terminus (red). For clarity, predicted unstructured residues are not shown (see S2 Fig, S1, S2 Tables for details). Cysteine pairs predicted to be within bonding distance either with high/medium probability or between chains have been labelled for one chain. Concentrated regions of cysteines that might participate in crystalline cross linking are indicated by X. Chain B, light grey; Chain C, dark grey. **B** Representation as above for the predicted CsxA timer.

A calculated map of a lattice of predicted trimers at 20 Å resolution was accommodated within the experimental density, providing a plausible molecular explanation for the observed architecture (Fig 4, S2 Movie). The fit accounts simultaneously for the crown-shaped sixfold assembly, the threefold interconnecting linkers, the dimensions of the unit cell, and the central pore without requiring major rearrangement of the predicted fold. The density that is not accounted for in the model fit is likely to represent parts of the chain that lie outside the envelope but that are predicted with low confidence; these include the loops pointing into the central pore. It should also be noted that density is likely to be smeared orthogonally to the crystal plane, given the lower resolution in this direction.

We previously observed a hydration state-dependent and reversible conformational reorganisation in the exosporium (Janganan 2020). In this work we compared hydrated and dehydrated crystals by AFM. The overall lattice dimensions and symmetry were retained following dehydration, but the surface topography surrounding the central pore changed markedly, indicating that the hydration-dependent conformational change is concentrated within the crown region. Gaussian-filtered representations of the cryoEM reconstruction closely reproduce the surface topography observed by AFM, allowing the two imaging modalities to be directly correlated. Comparison with the previously assigned AFM surface therefore allows the corresponding face of the cryoEM reconstruction to be assigned as the externally exposed surface of the exosporium (Fig 4, 6).

#### CsxB contributes primarily to envelope morphogenesis

Deletion of *csxB* produced localised blebbing of the outer coat. The exosporium remained intact and the parasporal layers were retained, indicating that CsxB is not a principal structural constituent of either assembly (Fig 3C, G, K). Thus, CsxB is not essential for formation of either the exosporium or the parasporal layers but appears to be essential for normal envelope morphogenesis.

#### CsxC is required for formation of the parasporal layers within the spore envelope

Deletion of *csxC* left the exosporium intact but completely eliminated the parasporal layers within the interspace (Fig 3D, H, L). The loss of these structures was accompanied by marked alterations in overall envelope morphology. The exosporium became tightly apposed to the coat around much of the spore body circumference, while remaining elongated at one pole (Fig 3D, H, 5B). In addition, the outer coat appeared frayed in all spores examined (Fig 3H, L). Thus, CsxC is essential for assembly of the parasporal layers while also indicating a broader role in envelope morphogenesis.

To define the organisation of the parasporal layers in greater detail, we examined the exposed layers in Δ*csxA* spores by AFM together with thin-section electron microscopy. Extended sheets were flattened out along with appendages (Janganan et al., 2016) wrapped round the spore coat and extending out from the spore body (Fig 7A). This shows that the exosporium is not required for the anchoring of one or more types of appendage. There are some appendages that appear to emerge from beneath the parasporal layers (Fig 7C). Therefore, it is possible that these are anchored deeper within the spore than the parasporal layers. The parasporal sheets possessed a well-defined crystalline lattice that is clearly distinct from the hexagonal symmetry of the CsxA exosporium basal layer (Fig 7B, E, G). Rather than forming a continuous single-layered sheet, the material occurred as multiple closely apposed laminae, consistent with the multilayered structures observed in thin-section electron microscopy of wild-type spores (Fig 2, 7C-G). In many regions the crystalline sheets were partially covered by disordered material (Fig 7F), consistent with the amorphous fringes apparent in electron micrographs of intact spores, suggesting that the two observations correspond to the same structural feature (Fig 1B-C).

**Fig 6.**
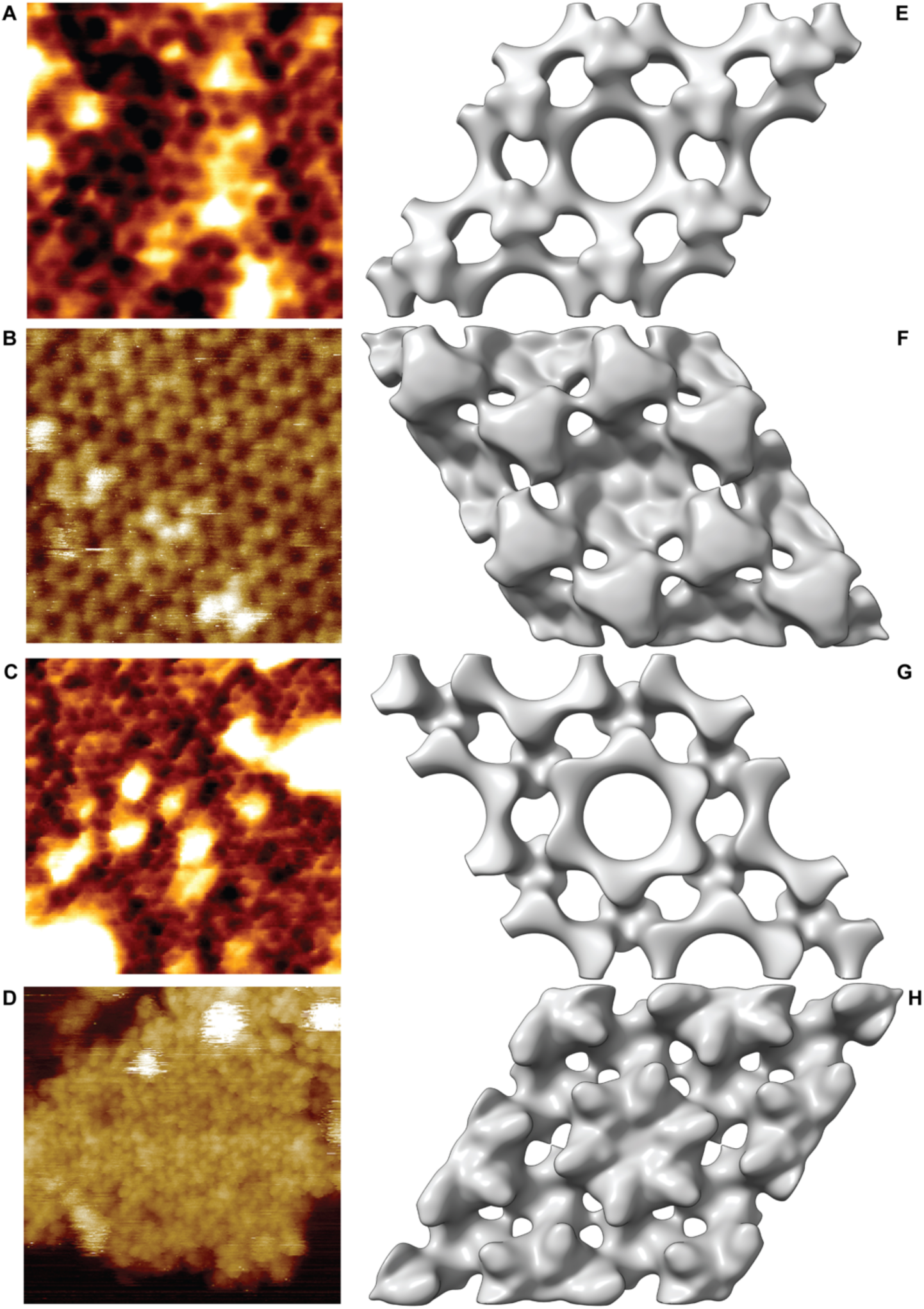
Comparison of hydrated cryoEM and AFM structures identifies the orientation relative to the spore core, and hydration-dependent conformational changes of the CsxA lattice. **A-D** AFM of recombinant CsxA crystals from Janganan *et al*. (Janganan et al., 2020). **E-H** CryoEM reconstructions of samples cognate to (A-D). **A** Height image of a recombinant dehydrated CsxA 2D crystal showing a honeycomb-like lattice with a unit cell of ∼102 Å. Color scale, 2.5 nm. **B** Height image of recombinant hydrated CsxA 2D crystal, showing a honeycomb-like lattice of ∼103 Å. Colour scale, 6 nm. **C** Height image of a recombinant dehydrated CsxA 2D crystal showing a lattice with an apparent spacing of ∼50. Colour scale, 2.5 nm. **D** Height image of recombinant hydrated CsxA 2D crystal fragment showing a hexagonal lattice of flower-like structures with a unit cell of ∼100 Å. Color scale, 15 nm. **E** EM reconstruction of negatively stained CsxA 2D crystal from Janaganan *et al*. (Janganan et al., 2020) showing a similar honeycomb structure to (A). **F** CryoEM reconstruction of hydrated CsxA from the current study (Fig 8). We interpret the prominent peaks in (B) to correspond to the prominent threefold linkers arrangement around a sixfold symmetric cavity leading into the pore. **G** EM reconstruction of negatively stained CsxA 2D crystal from Janaganan *et al*. (Janganan et al., 2020) showing the opposite face from (E). Panel (C) appears to show a unit cell of only ∼50 Å which we interpret to be the spacing between large and small pores revealed by EM. **H** View of face opposite to (F). We interpret the prominent ridges radiating from the central sixfold symmetry axis to correspond to the elevated ‘petals’ seen in (D). Since the view in (D) was assigned as the outward facing side by Janganan *et al*. (Janganan et al., 2020) we tentatively assign the view in (H) also as the outward face.

**Fig 7.**
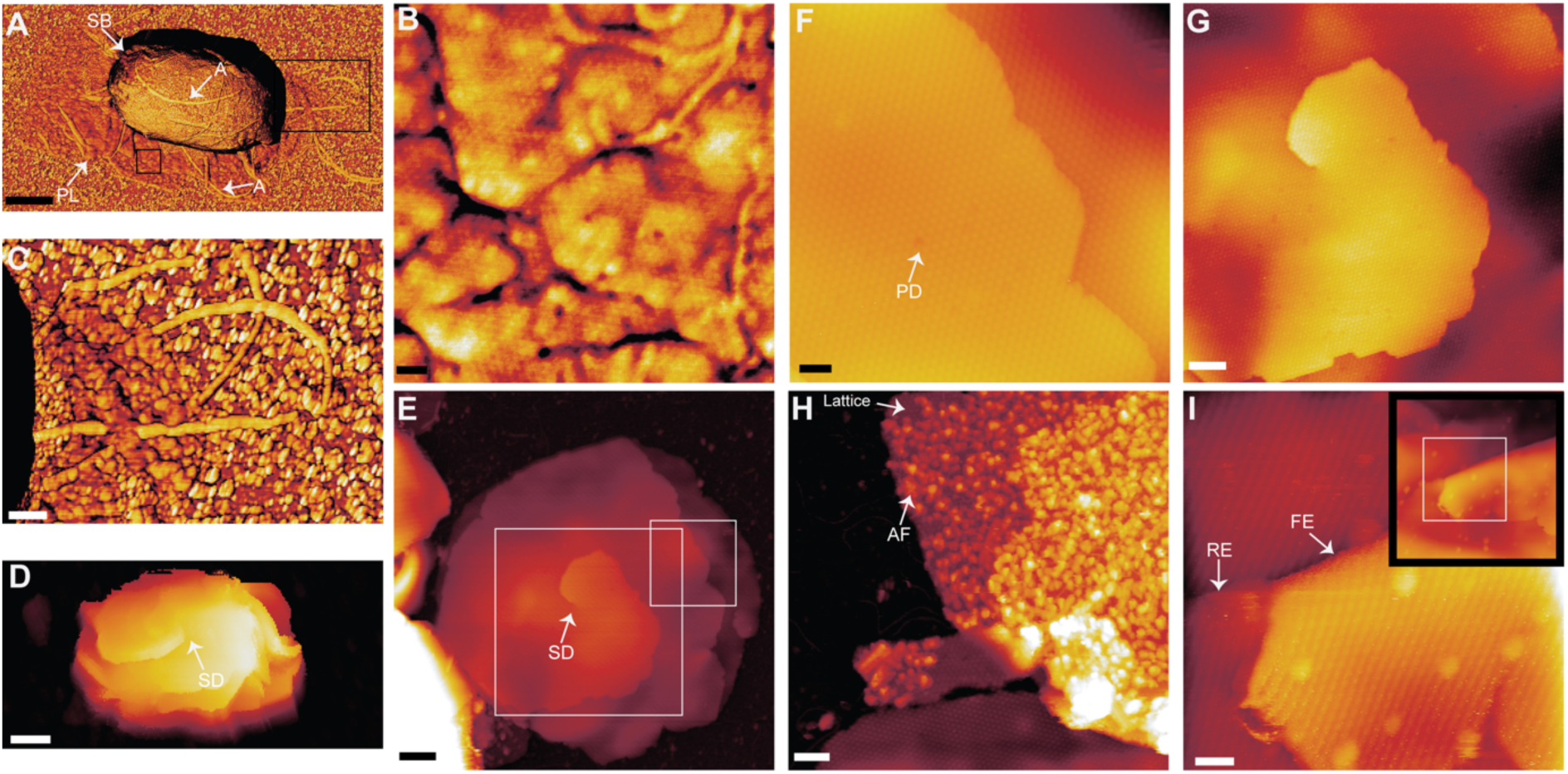
Parasporal layers form a three-dimensional crystalline assembly through screw-dislocation-mediated crystal growth. **A** AFM (phase image) of a dormant Δ*csxA* spore in air. Appendages (arrows, A) are visible on the spore body (SB) and its surroundings. The edges of the parasporal layers are observed surrounding the spore body and spreading on to the substrate (PL). Phase range: 30 degrees. Scale bar: 500 nm. **B** Higher magnification of (A) on the flat outer parasporal layers, this area displays the same lattice observed on isolated fragments (F). Phase range: 13 degrees. Scale bar: 20 nm. **C** Higher magnification of (A) showing appendages that emerge from beneath the flat outer parasporal layers. **D** AFM (height image) of dormant Δ*csxA* spore, in liquid. A screw dislocation (SD) is visible on the surface of the hydrated spore (arrow, SD). Height range: 0 - 1.5 μm. Scale bar: 500 nm. **E** AFM (height image) of a fragment of parasporal layer, in liquid, isolated from Δ*csxA* spores. Screw dislocations are visible in some sheets of crystal (SD). Height scale: 0 - 70 nm. Scale bar: 100 nm. **F** Higher magnification of the screw dislocation in (E) with a visible lattice and helicoid. Height scale: 0 - 25 nm. Scale bar: 50 nm. **G** Higher magnification image of (E), the crystal lattice is visible with several point defects (PD). Lattices from different layers appear to be in register. Height scale: 0 - 60 nm. Scale bar: 20 nm. **H** Sheets of crystal with disordered material covering some sections. This disordered material may be correspond to the amorphous fringe (AF) visible in thin section EM (Fig 1B-C). Height scale: 30 nm. Scale bar: 50 nm. **I** A highpass filtered image of a folded edge (FE) of a sheet of crystal showing the same lattice on either side, with inset of a larger scan area (without any post processing for flattening). The additional label shows the rough edge (RE). Height scale: 45 nm, 100 nm (inset). Scale bar: 20 nm. Images A - C, and E-I collected in tapping mode, D collected in QI mode.

Thus, the parasporal layers are not derived from fragmentation of the exosporium but represent a genetically and structurally distinct component of the spore envelope.

#### Parasporal layers form via screw-dislocation-mediated crystallisation

AFM images of fragments of isolated parasporal layers showed a peak-to-peak lattice spacing of 63 ± 6 Å, (n = 31) but with numerous screw dislocations and point defects within the otherwise intact crystalline sheets (Fig 7C-D). The crystal lattice could be resolved continuously around the dislocation cores (Fig 7F), demonstrating that these features are an intrinsic feature of a coherent crystal rather than tears or folds generated during specimen preparation. The multiple layers of these fragments were stacked with their lattices in register (Fig 7D-E), with step heights between layers of 50 to 70 Å, n = 18. In some cases we could also image these same structures on the exposed surface of intact spores (Fig 7C).

#### CryoEM confirms a trigonal lattice in parasporal crystals

Since the parasporal layers formed stacks of variable numbers of layers, they were not accessible to three-dimensional structure determination by electron crystallography. Instead, we adopted a single particle approach, aligning images of crystal patches to generate 2D class averages (Fig 8A). We selected class (iii) for further analysis as it showed clear crystalline symmetry elements (Fig 8B). This averaged projection view has an apparent unit cell of *a* ≈ *b* ≈ 65 Å, γ ≈ 120°, consistent with AFM measurements above and consistent with the ∼50 Å ≈ repeat observed in S2 Fig (∼55 Å lattice spacing would be expected for a 65 Å unit cell). The projection is also consistent with *p*312 two-sided plane group symmetry where both sides should be equivalent (Holser, 1958; Amos et al., 1982); indeed AFM views of both faces of folded layers show both surfaces to be the same (Fig 7G). The close agreement between lattice dimensions determined independently by AFM, thin-section EM and cryoEM confirms that all three approaches visualise the same crystalline assembly.

**Figure 8.**
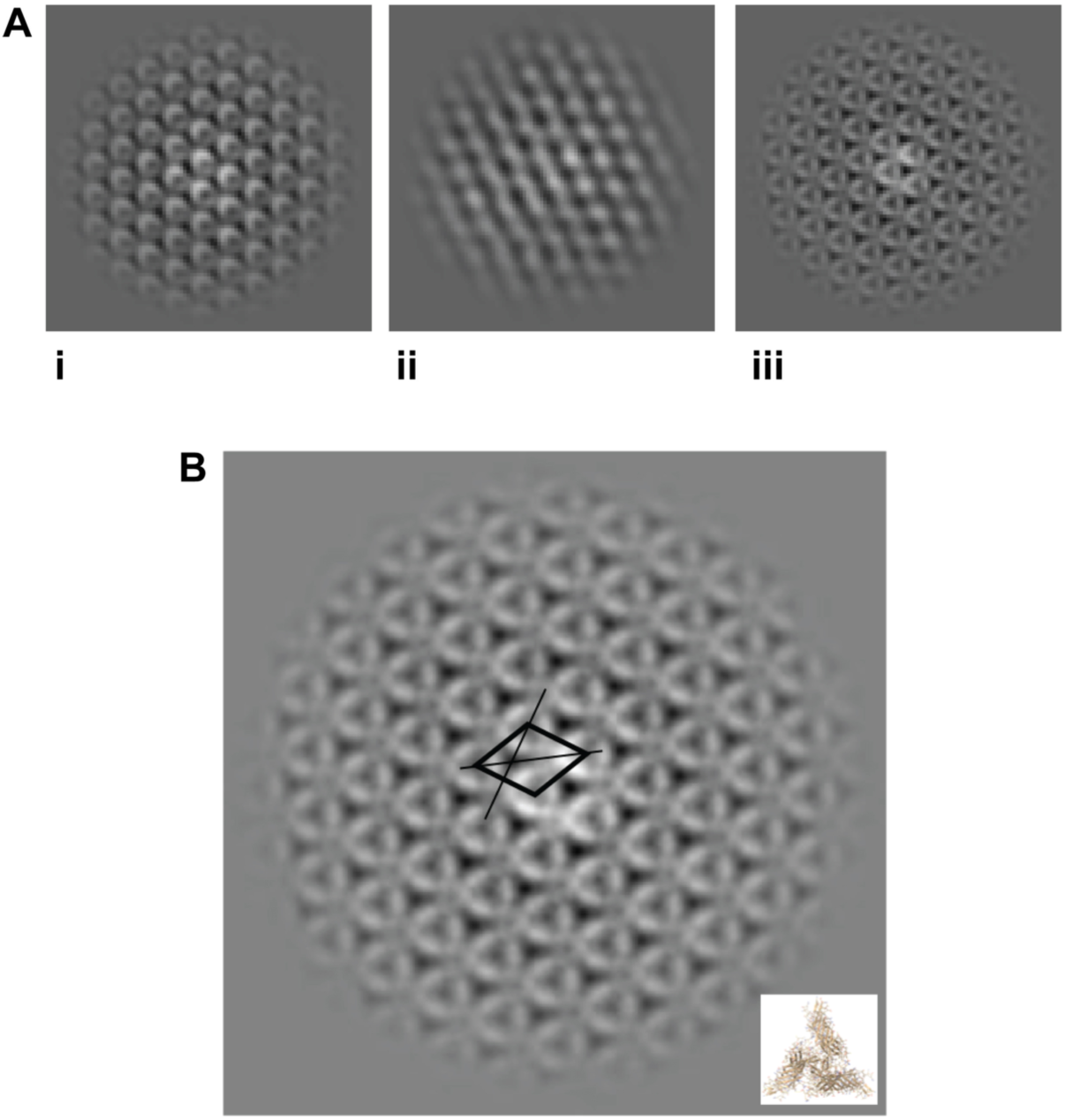
CryoEM independently confirms the trigonal architecture of the parasporal lattice. **A** Three class averages generated from ∼47,000 crystal patches. **B** Higher magnification view of class average (iii). The unit cell is shown by the thick solid lines (*a* ≈ *b* ≈ 65 Å, *γ* = 120°). Two in-plane twofold symmetry axes have been shown by thin solid lines. Inset shows a projection of a putative CsxC trimer for comparative scale.

#### Structural modelling and mass spectrometry support CsxC as the principal component of the parasporal layer

Mass spectrometry analysis of spore surface layers isolated from Δ*csxA* spores identified the 3 most abundant proteins, and most likely candidate proteins for parasporal layer composition, as CsxB (38%), an uncharacterised hypothetical protein (locus tag: 03300)(7.3%), and CsxC (7.2%)(S1 Data). Despite the high abundance of CsxB it is an unlikely structural candidate as Δ*csxB* spores retain their parasporal layers (Fig 3G, K). Δ*csxC* spores on the other hand do not contain parasporal layers making CsxC the most likely structural candidate for parasporal layer composition.

To determine whether the observed lattice organisation is consistent with the properties of CsxC, we modelled the protein using AlphaFold3 (Abramson et al., 2024). As for CsxA, CsxC is predicted to assemble as a trimer (Fig 5) with the threefold symmetry compatible with the trigonal parasporal lattice observed experimentally. The predicted trimer forms a tripod-like arrangement with each leg composed of a β-barrel made up of two three-stranded sheets, characteristic of the SPOCS domain as described for CsxA above. The C-terminal proximal sheet of chain A is extended by two long additional β-strands from chain B and a short strand from chain C (Fig 5A, S2 Fig). Each subunit contains 14 cysteine residues, several of which are positioned in the AlphaFold3 model such that disulphide bonds could feasibly form. (Fig 5A, S2 Fig and S2 Table). Such cross-linking could contribute to stabilisation of the crystalline assembly (Fig 7A, S2 Fig, S2 Table).

We have not attempted to interpret the projection map by modelling the molecular packing because with *p*312 symmetry there may be superimposition of trimers with ‘up’ and ‘down’ orientations, making the packing impossible to interpret from one projection view; we previously reported an analogous overlapped packing arrangement for *B. subtilis* CotY assemblies where interpretation of the crystal packing only becomes possible with a full 3D analysis (Jiang et al., 2015).

#### BclA and BclB are predicted collagen-like envelope proteins

In Δ*bclA* spores the crystalline exosporium remained intact, whereas the external filamentous layer, known as the hairy nap, was substantially altered. Although the longer (∼100 nm) filaments were absent, a shorter (∼30 nm), disordered layer remained associated with the outer surface of the exosporium (Fig 9D). BclA is expected to adopt the characteristic organisation of trimeric bacterial collagen-like proteins (Sylvestre et al., 2003; Réty et al., 2005), comprising an extended collagenous stalk. AlphaFold3 predicts a globular N-terminal domain (S2 Fig) which adopts a similar fold to that from the crystal structure of the *Bacillus anthracis* BclA (PDB ID: W1CK, 3TYJ)(Réty et al., 2005; Kirchdoerfer et al., 2012). Based on sequence BclA is predicted to be approximately 150 nm in length and BclB approximately 30 nm. BclB does not have any close structural homologues in the PDB but has a similar predicted fold of its C-terminal domain as BclB of *B. anthracis* (S2 Fig); it is predicted to have a collagen-like N-terminal tail of ∼30 nm length. Deletion of *bclB* produced no detectable alteration in spore morphology. The exosporium, parasporal layers and hairy nap remained indistinguishable from those of wild-type spores, suggesting that BclB is not a major structural component of the mature dormant envelope (Fig 9C). However, if BclB is normally masked by the longer BclA, we would not expect to detect its absence in this mutant.

**Figure 9.**
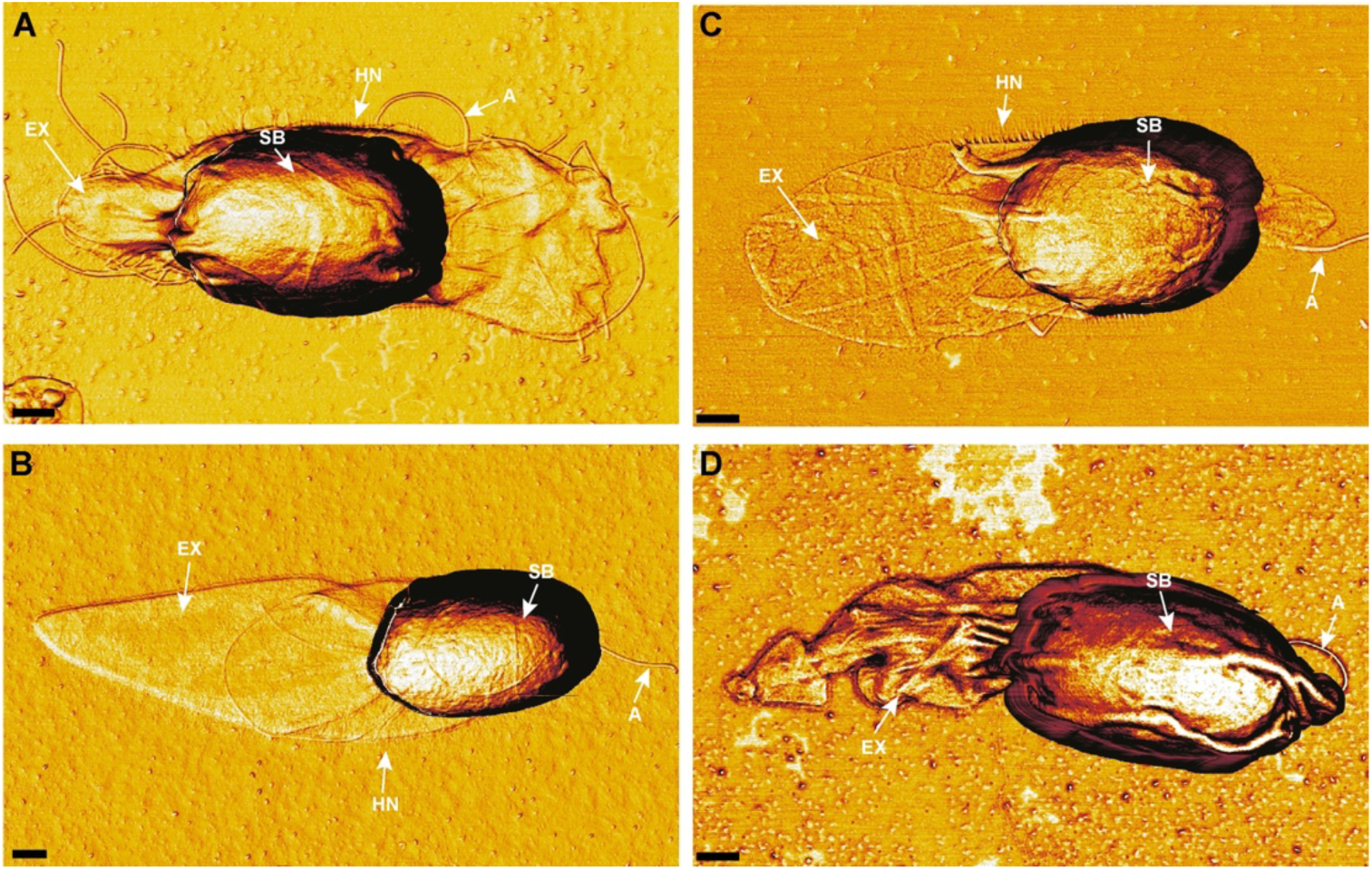
Topographical differences of dormant spores from deletion mutants by AFM. AFM phase images of dried dormant spores in air. Notable labelled features include the exosporium (EX), the hairy nap (HN), appendages (A), and the spore body (SB). **A** and **B** Both *ΔcsxB* and *ΔbclB* maintain all salient topographical features of WT spores listed above and appear the same as WT spores (Fig 1E). **C** *ΔcsxC* also maintains these features but the exosporium appears more tightly apposed to the spore body with a highly extended exosporium. **D** *ΔbclA* lacks the appearance of the hairy nap fringe that extends approximately 100 nm from the exosporium. All other features remain comparable with WT spores. Scale bar: 250 nm. Phase ranges: **A** 23°, **B** 17.8°, **C** 23.6°, **D** 12.2°. N = 10 spores.

### Effect of mutations on sporulation and environmental interactions

In addition to interrogating the architectural roles of the envelope proteins we also examined whether deletion of these proteins affected sporulation. Deletion of *csxB*, *csxC*, *bclA* or *bclB* had no detectable effect on sporulation efficiency (S3 Fig). In contrast, deletion of *csxA* resulted in a small reduction in efficiency.

Mature spores are extremely resilient to a range of environmental insults. To determine if envelope mutations impacted heat resistance, purified spore suspensions were incubated at 65 °C, the temperature used in the sporulation assay, or at 75, 85 and 95 °C. Wild type spores were unaffected by incubation at 75 °C, but 85 °C saw an ∼50-fold drop in viability. Spore viability dropped below the level of detection at 95 °C (S4 Fig). In general, mutant strains showed no difference in spore viability after treatment at 65 °C and 75 °C and a similar drop compared to WT at 85 °C. Thus, despite producing pronounced changes in outer-envelope architecture, deletion of these individual components had little effect on the intrinsic heat resistance of mature spores.

### Hydration-dehydration cycling of wild-type and mutant spores

We next examined the response to the spore envelope to sequential cycles of dehydration and rehydration. Wild-type spores underwent pronounced and only partially reversible structural changes in response to hydration state. Dehydration caused the exosporium to collapse tightly around the spore body and onto the underlying substrate, producing a characteristic “shrink-wrapped” morphology (Fig 1G-H). This was accompanied by an approximately fourfold reduction in measured spore volume (S5A Fig). Following rehydration, spores recovered to approximately half of their original hydrated volume. Although parts of the exosporium re-expanded, regions, predominantly the edges of the extended exosporium, remained locally collapsed against the substrate (S5G Fig).

Spores lacking the exosporium (Δ*csxA*) exhibited hydrated and dehydrated volumes similar to those of wild-type spores (S5A Fig). However, the volume of hydrated Δ*csxA* spores may be overestimated because the exposed parasporal layers adopt a splayed morphology, projecting outwards from the spore surface (Fig 4C, S5H Fig). This arrangement is consistent with thin-section electron micrographs, which show separation between parasporal layers, particularly at the spore poles (Fig 3F, J). Because the AFM tip cannot access the space beneath these projecting layers, the volume reconstruction effectively treats the region below the maximum detected height as occupied, potentially inflating the measured volume. Following dehydration and subsequent rehydration, the parasporal layers did not fully recover their initial splayed arrangement (Fig S5J), which may therefore account, at least in part, for the incomplete recovery of the measured spore volume.

## Discussion

The sporulation programme is deeply conserved across the Bacillota, yet proteins forming the spore’s outermost protective structures are highly divergent. The *C. sporogenes* spore envelope shows that similar architectures can arise from structurally distinct proteins through common self-assembly principles. Unexpectedly, one assembly exploits screw-dislocation mediated crystal growth to generate a multilayered three-dimensional crystal-a mechanism rarely described in native biological assemblies.

### An integrated model of the *C. sporogenes* spore envelope architecture

Our data support a hierarchical model for assembly of the *C. sporogenes* spore envelope (Fig 10). CsxA forms the two-dimensional crystalline exosporium basal layer, decorated with BclA which generates the hairy nap. The exosporium encloses the interspace within which CsxC forms the parasporal layers, separating the exosporium and coat. CsxB contributes to coat and envelope organisation. The makeup of the interspace is poorly characterised, but it appears to be a hydrated and dynamic compartment containing proteins and likely also polysaccharide (Lehmann et al., 2022). None of the proteins characterised here accounts for the additional appendages observed in mature spores, other than the hairy nap (Fig 1E, 7A, 9), indicating that additional components remain unidentified. Whether these structures are anchored deeper within the spore envelope or some originate within the exosporium also remains unknown.

**Figure 10.**
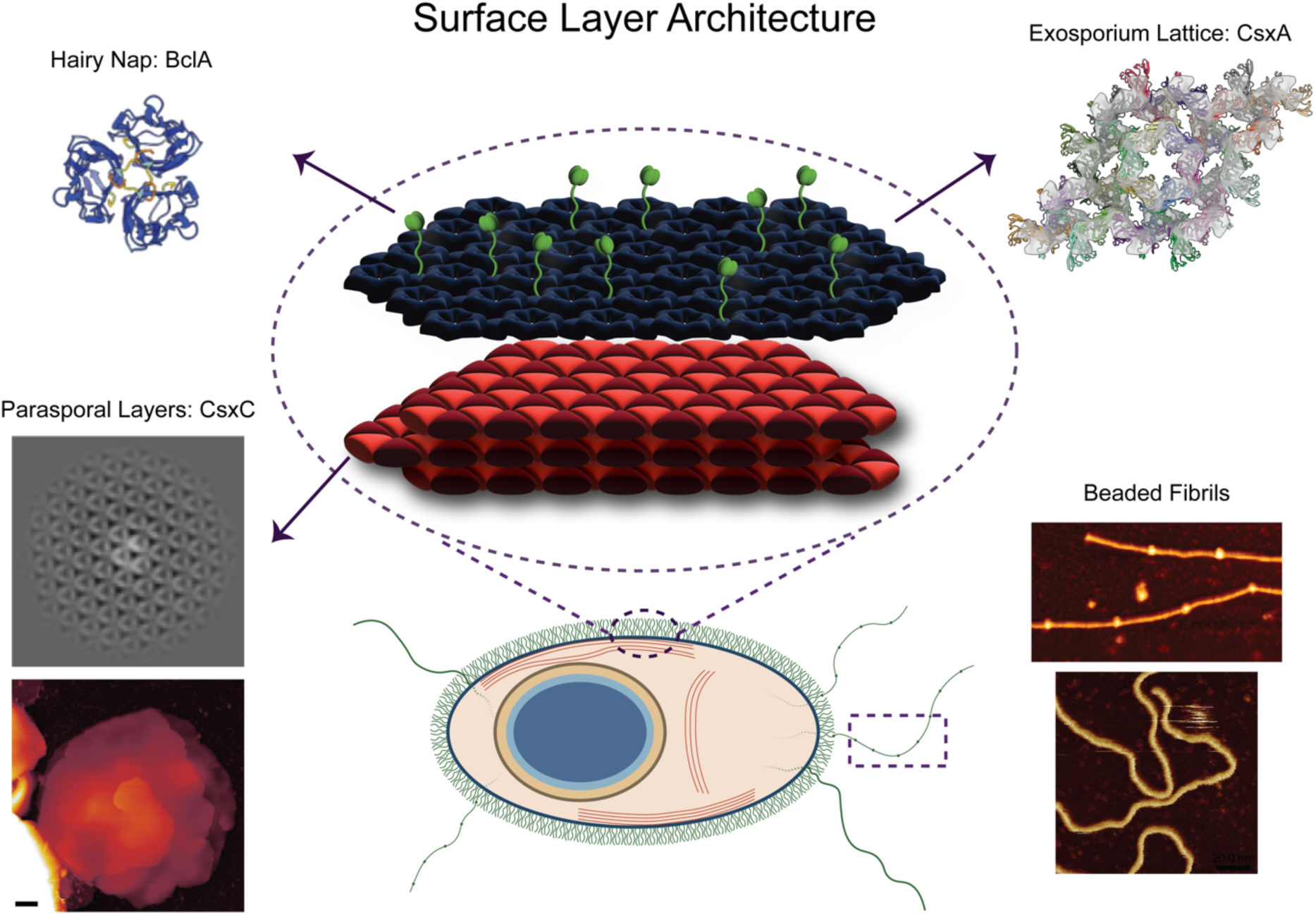
Integrated molecular model of the *Clostridium sporogenes* spore envelope. Schematic representation incorporating the genetic and structural observations from this study. The crystalline exosporium basal layer is formed by CsxA and is decorated externally by the BclA-containing hairy nap. The interspace between the exosporium and coat contains the crystalline parasporal layer assemblies whose formation requires CsxC; CsxC is shown as its proposed principal structural component. CsxB is required for normal envelope morphogenesis but has not been assigned to a specific structural compartment. The specialised polar region is shown schematically; its molecular composition and mechanism of opening remain unknown. Various types of appendage extend out from the external surface of the spore; we have only represented the beaded fibrils in hydrated and dehydrated forms, as we previously described (Janganan 2016).

At one pole of the exosporium we observed structural differences, either an aperture or lipped cap. The aperture (Fig 1D, G, H) is potentially sealed in dormant spores by the mechanically fragile cap as observed by others (Panessawarren et al., 1994; Brunt et al., 2015). If so, vegetative cell emergence may depend on local mechanical failure at this pole rather than enzymatic digestion of the exosporium (Hoeniger and Headley, 1969).

The outer envelope thus comprises two characterised and complementary protein classes: cysteine-rich DUF3794/SPOCS domain CsxA and CsxC forming robust crystalline scaffolds, and collagen-like proteins decorating the external surface. The following sections consider how these structures are assembled to meet their functional requirements.

### Distinct crystalline arrays formed by cysteine-rich CsxA and CsxC

Regular hexagonal-like arrays were first recognised in Clostridial outer envelopes five decades ago (Masuda et al., 1980; Takumi 1979; Mackey 1972). Here we confirm that for *C. sporogenes* CsxA forms the robust exosporium basal layer consisting of a highly interlinked network of protein subunits (Fig 4). The absence of the exosporium in Δ*csxA* spores identifies this layer as a genetically distinct structural module (Fig 3). Mutant spores retain an intact cortex, coat, interspace and parasporal layers, demonstrating that the basal layer is dispensable for assembly of the core spore body (Fig 3, 7A).

The AlphaFold3 prediction of CsxA as a trimeric tripod-like complex plausibly fits the cryoEM density of recombinant CsxA crystals to recapitulate the native basal layer (Fig 4C-D, S2 Movie)(Janganan et al., 2020). The model positions several of the 25 cysteine residues in each CsxA subunit sufficiently close to form potential intra- and intermolecular disulphide bonds (Fig 5B, S2 Fig, S1 Table), consistent with evidence that the array is reinforced by reducible cross-links (Fig 4) (Janganan et al., 2020). Several cysteines in the model are unpaired, notably at each tripod leg (Fig 5B) - these may cross-link adjacent trimers. These putative bonds provide a testable explanation for the extreme resistance of CsxA crystals to chemical and thermal denaturation and parallel other cysteine-rich spore proteins that exploit cooperative disulphide cross-linking to form robust assemblies (Jiang et al., 2015; Terry et al., 2017)(Romero-Rodríguez et al., 2020; Janganan et al., 2020).

The extensive cross-linking must nevertheless be compatible with structural flexibility. The pronounced reduction in spore volume during dehydration and ‘shrink-wrapped’ exosporium demonstrate that it is dynamic rather than rigid. Individual CsxA subunits undergo reversible structural changes during hydration cycles (Janganan et al., 2020). However, comparison of hydrated cryoEM and dried, negatively stained structures (Janganan et al., 2020) indicates that these rearrangements involve local changes in pore geometry rather than large-scale lattice reorganisation (Fig 6).

CsxC is responsible for a second and fundamentally different crystalline system within the same envelope. Lamellar material beneath the exosporium had been described previously in *C. sporogenes* (Hoeniger and Headley, 1969) and other Clostridia (Lund et al., 1978; Masuda et al., 1980; Portinha et al., 2022); these also encode CsxC homologues, suggesting a common molecular basis. The parasporal layers therefore represent a genetically distinct assembly rather than an elaboration of the exosporium (Fig 1B, C, 3, 7).

Several independent observations identify CsxC as the principal component of these parasporal layers. The loss of parasporal layers in Δ*csxC* spores supports CsxC as a principal structural constituent rather than solely a morphogenetic factor. Mass spectrometry shows that CsxC is one of the most abundant species in the envelope material of Δ*csxA* spores. Moreover, isolated sheets exposed in Δ*csxA* spores exhibit an apparently trigonal crystalline lattice distinct from the hexagonal CsxA lattice, consistent with a different self-assembling protein (Fig 2, 7, 8). AlphaFold3 predicts a trimeric CsxC assembly compatible with the experimentally observed threefold symmetry (Fig 5, S7 Fig). The structural similarity and cysteine-rich content of CsxA and CsxC further suggest that CsxC also self-assembles to form a robust crystalline lattice reinforced by disulphide bonds (Fig 5).

Δ*csxC* spores also display defects beyond loss of parasporal layers. The exosporium becomes abnormally elongated at one pole while wrapping tightly around the remainder of the spore, and the underlying coat frequently appears frayed (Fig 3). Thus, CsxC contributes to outer-coat organisation and influences final exosporium architecture, serving both as a structural component of the parasporal layers and a morphogenetic determinant of neighbouring envelope layers.

Observations under varying hydration states are consistent with a mechanical role for CsxC. The similar dehydrated volumes of wild-type and Δ*csxA* spores indicate that the exosporium contributes little to dehydrated spore dimensions. In contrast, comparison of wild-type with the smaller dehydrated Δ*csxC* spores (S5A Fig) suggest that parasporal layers contribute substantially to envelope bulk. The Δ*csxA* mutant indicates that parasporal layers are also hydration-sensitive, although they do not collapse like the exosporium, reflecting their distinct construction. Their splayed morphology was reduced following dehydration and rehydration (S6H-J Fig), potentially reflecting altered higher-order organisation rather than irreversible changes to individual layers The tendency of the to remain collapsed in Δ*csxC* spores suggests that parasporal layers maintain separation between opposing exosporium regions during hydration cycling. Without them, increased contact during dehydration may restrict subsequent re-expansion.

Together, the CsxA basal layer and the CsxC parasporal layers establish that the Clostridial spore envelope is constructed from at least two genetically independent crystalline systems with distinct architectures, mechanical properties and specialised functions.

CsxB, which appears to be a highly abundant protein, occupies a different position in the envelope hierarchy. Deletion of *csxB* produces only subtle structural defects, including outer-coat blebs and altered coat-exosporium spacing (Fig 3C, G, K, 9A). CsxB is predicted to adopt a von Willebrand factor A-like fold associated with protein–protein interactions, distinct from CsxA and CsxC (S2 Fig), suggesting an assembly role rather than a principal structural function.

### Parasporal layers assemble by a biologically unusual mode of three-dimensional crystal growth

Assembly of the parasporal layers differs fundamentally from that of CsxA or the cysteine-rich exosporium proteins ExsY and CotY of *Bacillus anthracis/cereus/thuringinesis* (Terry et al., 2017; Janganan et al., 2020). Whereas CsxA, ExsY and CotY form continuous crystalline sheets one molecule thick, the putative parasporal protein CsxC produces multilayered three-dimensional assemblies whose successive layers remain crystallographically aligned (Fig 7D-G). These layers are interconnected by screw dislocations, supporting three-dimensional growth mediated through these screw dislocations rather than through repeated nucleation of independent sheets; this a kinetically favourable mechanism for constructing these multilamellar assemblies.

Although screw dislocations are well recognised in inorganic and artificial protein crystals (Frank, 1949; Pum et al., 2021; Reviakine et al., 2003; Malkin et al., 1995), examples in native biological protein assemblies are exceptionally rare. Previous studies of protein layers within spores of *Clostridium novyi* by AFM (Plomp et al., 2007) and *Clostridium pasteurianum* by thin section EM (Mackey and Morris, 1972) inferred screw or edge dislocations but lacked sufficient resolution to establish a surrounding crystalline lattice. Here, we directly resolve the lattice around dislocation cores, confirming that they are genuine crystalline defects (Fig 2, 7D-G). Screw-dislocation-mediated growth has been observed in biomineralised shell structures, including nacre (mother-of-pearl), our observations now suggest that this classical crystal-growth mechanism can be exploited in both mineralised and proteinaceous biological materials (Yao et al., 2006).

Thus, a native cellular structure appears to exploit a fundamental crystal-growth mechanism for biological morphogenesis. We propose that parasporal-layer robustness derives from two complementary features: extensive disulphide cross-linking within individual sheets and their integration into a multilamellar crystal through screw-dislocation-mediated growth.

### CsxA and CsxC share structural features with morphogenetic and structural proteins of other spore formers

AlphaFold3 (Abramson et al., 2024) predicts that CsxA and CsxC share the SPOCS/DUF3794 fold found in several morphogenetic proteins from other Bacillota, indicating structural homology between proteins with distinct functions (Fig 5, S2 Fig)(Delerue 2022). Their tripod-like assemblies closely resemble that predicted for the *Bacillus subtilis* morphogenetic protein CotE (partially confirmed by cryoEM), despite the absence of sequence similarity (S6 Fig)(Lee et al., 2025). In all the three proteins, CsxA, CsxC and CotE, interlocking β-strands between neighbouring subunits create a stable trimeric hub, suggesting a conserved structural solution for spore protein scaffolds.

Despite this common core, the proteins form distinct supramolecular architectures. Unlike CotE, CsxA and CsxC contribute additional β-strands to partner subunits within their β-barrel domains (S7 Fig). CotE forms open mesh-like three-dimensional arrays (Aronson et al., 1992; Jiang et al., 2015; Lee et al., 2025) and acts primarily as an outer-coat morphogenetic organiser (Zheng et al., 1988; McKenney et al., 2013; Bauer et al., 1999; Little and Driks, 2001; Driks, 1999; Henriques and Moran, 2000; Henriques and Jr., 2007; Bauda et al., 2024), whereas CsxA forms the two-dimensional exosporium basal layer and CsxC assembles into densely packed three-dimensional crystals in the interspace. Their higher cysteine content is likely to enable extensive disulphide cross-linking and account for these different assembly modes. Thus, a shared ancestral fold appears to have diversified for distinct architectural functions, supporting DUF3794/SPOCS proteins as a conserved scaffold adapted to different roles during spore envelope evolution (Delerue et al., 2022).

AlphaFold predictions also indicate that cysteine-rich proteins implicated in spore envelope formation in *C. difficile* (CdeC) (Romero-Rodríguez et al., 2020) and *C. sordellii* (CsA and CsB) (Rabi et al., 2017; Rabi et al., 2018) adopt related folds (S7 Fig). In particular, *C. sordellii* CsA and CsB behave similarly to *C. sporogenes* envelope proteins, with both required for normal spore architecture. CsA shares 42% identity with the *C. difficile* exosporium protein CdeC (S7 Fig), reported as a major exosporium component (Barra-Carrasco et al., 2013); however, *C. difficile* spores lack the ‘sac-like’ exosporium of *C. sporogenes* and other Bacillota. CdeC is tightly integrated into the exosporium and may form an internal scaffold supporting the external CdeM layer (Antunes et al., 2018).

The *C. difficile* morphogenetic protein SipL (Putnam et al., 2013; Touchette et al., 2019) adopts a similar tripod architecture but within a single polypeptide. SipL functions as an envelope assembly factor, and AlphaFold3 predicts three tandem DUF3794 domains with pseudo-threefold symmetry (S7 Fig), closely resembling the intertwined CsxA and CsxC trimers. This further supports a common ancestral scaffold for morphogenetic and structural envelope proteins. Like CotE, however, SipL lacks cysteines positioned to form disulphide bonds.

Conservation of this architecture across diverse Clostridia expands DUF3794/SPOCS proteins into a broad family of spore structural and morphogenetic proteins. An ancestral fold may have diversified into two classes: cysteine-rich proteins, including CsxA, CsxC, CdeC and CsB, that construct tightly packed, cross-linked arrays, and cysteine-poor proteins, including SipL and CotE, that primarily perform morphogenetic organising roles.

### Cysteine-rich spore envelope proteins follow a conserved physico-chemical strategy

We predict that CsxA and CsxC form extensive networks of intra- and intermolecular disulphide bonds (Fig 4, 5 S6 Fig). Together with evidence that the *Bacillus* proteins CotY and ExsY self-assemble into stable two-dimensional crystals, these findings support cysteine-rich, disulphide-stabilised self-assembly as a fundamental principle of spore envelope construction (Jiang et al., 2015; Terry et al., 2017; Janganan et al., 2020; Romero-Rodríguez et al., 2020).

The analogy extends to the porous basal-layer templates bearing collagenous hairy nap proteins found for both CsxA in *C. sporogenes* and ExsY arrays in *B. cereus*. Although it remains unclear whether the *C. sporogenes* exosporium is completely sealed (Fig 1D-H), the central pores of the CsxA lattice are approximately 25 Å at their narrowest. This is smaller than many globular enzymes and antibodies, consistent with a molecular sieve that permits diffusion of small molecules, including water, required for germination and metabolic reactivation. This parallels the proposed sieving function of the ExsY basal layer in the *B. anthracis/cereus/thuringiensis* group (Ball et al., 2008; Kailas et al., 2011; Terry et al., 2017), suggesting selective permeability is conserved despite divergence of exosporium proteins.

The one exosporium protein apparently conserved in structure and function between *C. sporogenes* and the *B. cereus* group is BclA. As for ExsY, its predicted trimeric organisation suggests a symmetry match with the threefold features of the CsxA lattice, with attachment on its outward face (Fig 6, 10)(Terry 2017). Precise localisation will require analysis of intact nap-bearing exosporium or mapping interactions between the BclA C-terminal region and CsxA.

The physiological role of the exosporium in *C. sporogenes* remains to be established. Δ*csxA* spores remain viable and show only modest changes in sporulation efficiency and heat resistance (S3, S4 Fig), although the exosporium contributes to heat resistance in some *C. botulinum* strains (Portinha et al., 2022). Given its substantial metabolic cost in assembly, the *C. sporogenes* exosporium may instead be essential for environmental persistence, modulation of germination and outgrowth, or protection against stresses not reproduced in the laboratory.

### Have the exosporia of Bacillus and Clostridium evolved independently?

Core sporulation pathways are highly conserved throughout the Bacillota, indicating that endospore formation arose early in the phylum (Galperin et al., 2022). In contrast, few coat or exosporium proteins are shared between classes, consistent with our structural observations. Although the major *B. cereus* group exosporium proteins ExsY and CotY also form crystalline arrays, their predicted tertiary and quaternary structures differ markedly from CsxA and CsxC. ExsY and CotY form hexameric rings surrounding central pores (Terry 2017; Ball 2008), and their AlphaFold3-derived structures agree closely in morphology and dimensions with previous EM and AFM data while predicting a distinct α/β subunit fold (S7 Fig)(Ball et al., 2008; Terry et al., 2017; Kailas et al., 2011). Despite these differences, both systems generate highly ordered, resistant, disulphide-stabilised outer shells.

These findings suggest that Bacilli and Clostridia converged on similar solutions for constructing robust extracellular barriers. In both, symmetric protein assemblies reinforced by disulphide cross-linking generate stable protective layers despite distinct evolutionary origins. This represents convergent evolution at the level of supramolecular architecture, analogous to the independent emergence of crystalline S-layers across bacterial and archaeal phyla (Fagan and Fairweather, 2014; Johnston 2024; Barwinska-Sendra et al., 2025). Thus, evolution may conserve the physico-chemical principles of durable cellular envelopes while replacing the underlying molecular scaffold.

## Materials and Methods

### Strains and Growth Conditions

*C. sporogenes* NCIMB 701792 (NCDO 1792) was grown at 37 °C either in tryptone yeast (TY) broth or on brain heart infusion (BHI) agar in an anaerobic workstation (Don Whitley Scientific) with an atmosphere composed of 80% N_2_, 10% H_2_ and 10% CO_2_. *Escherichia coli* was grown in LB broth or on LB agar. *E. coli* strain NEB5*α* (New England Biolabs) was used for routine cloning and plasmid propagation. *E. coli* strain CA434 was used as a conjugation donor. Growth media were supplemented with chloramphenicol (15 µg ml^−1^), thiamphenicol (15 µg ml^−1^), colistin (50 µg ml^−1^), erythromycin (5 µg ml^−1^) or 4% (w/v) xylose as appropriate.

### Spore Preparation

A 5 ml overnight culture of *C. sporogenes* in TY broth was subcultured to an OD_600nm_ of 0.1 in 20 ml TY broth and incubated anaerobically for a minimum of 10 days. Cultures were harvested by centrifugation at 6,000 g for 5 min. Pelleted material was washed in 10 ml ice-cold sterile water 5 times before being resuspended in 0.5 ml 20% (w/v) HistoDenz (Sigma). This was layered atop 1 ml 50% (w/v) HistoDenz and centrifuged at 15,000 g for 15 min to separate spores from cell debris. HistoDenz was removed by aspiration and the spore pellet was washed a further 5 times in ice-cold sterile water to remove any residual HistoDenz. The purity of each preparation was checked by phase contrast microscopy.

### Sporulation Efficiency

Sporulation assays were performed based on methods developed for *Clostridioides difficile* (Dembek 2015). Overnight TY broth cultures of *C. sporogenes* were diluted to an OD_600nm_ of 0.01 in fresh pre-reduced broth and incubated for 12 h, and diluted again to an OD_600nm_ of 0.0001 before growth overnight to stationary phase. This minimised the carryover of spores from initial cultures. The relative proportions of vegetative cells and spores were then monitored at 24 h intervals for 5 days. At each time point, the total number of bacteria was determined by counting the number of CFU on TY agar. To determine the number of spores, samples were incubated at 65°C for 30 min, prior to CFU enumeration.

### Heat Resistance

*C. sporogenes* spore samples at an OD_600nm_ of 0.5 were incubated at RT, 65, 75, 85 and 95 °C for 30 min. Following serial dilution samples were plated onto TY agar for CFU enumeration to determine spore viability after heat treatment.

### Genome Sequencing

The genome of *C. sporogene*s strain NCIMB 701792 was sequenced to allow for precise manipulation of the genome. gDNA extraction, library prep (Nextera XT Library Prep Kit (Illumina, San Diego, United States of America) and Oxford Nanopore Technologies SQK-RBK114.96 kit (ONT, UK)), 30× illumina sequencing (NovaSeq 6000, 250-bp paired end protocol) and nanopore sequencing (GridION, FLO-MIN114 (R.10.4.1) flow cell) was performed at MicrobesNG (Birmingham, UK). Illumina reads were trimmed at MicrobesNG using Trimmomatic (v0.30) with a sliding window quality cutoff of Q15 (Bolger et al., 2014). Trimmed reads were checked using FastQC (v0.11.9) (https://www.bioinformatics.babraham.ac.uk/projects/fastqc/). *De novo* assembly was performed using Unicycler (v0.5.1)(Wick et al., 2017), and annotated with Prokka (v1.14.6)(Seemann, 2014).

### *C. sporogenes* Mutagenesis

Mutant strains of *C. sporogenes* NCIMB 701792 were constructed via allelic exchange mutagenesis. Mutagenesis plasmids contained either the pJAK184 backbone or slightly modified versions of pJAK184 (pJAK188 and pHF012, see S5 table). All of these are derivatives of pMTL-SC7215 (Cartman 2012) in which the *codA* gene has been replaced by a xylose inducible *mazF* (Fuchs et al., 2021). All strains, plasmids and primers can be found in S4-6 Tables.

Constructs for mutagenesis were produced as follows: for *csxB* deletion, homology arms complementary to regions immediately upstream and downstream of *csxB* were amplified by PCR (RF1686/1687, RF1688/1689 respectively) and inserted into PCR linearised pJAK184 (RF311/312) via Gibson assembly to produce pHF001. An erythromycin resistance cassette was constructed by cloning the *ermB* gene and *fdx* terminator from pRPF215 (RF1877/1878)(Dembek et al., 2015) downstream of the *cwp2* promoter in pJAK014 between the SacI and BamHI sites to produce pHF007 (Oatley et al., 2020; Fisher et al., 2026). The *ermB* cassette was then amplified (RF1886/1887) and inserted into PCR linearised pHF001 (RF1824/1825) via Gibson assembly to produce the final construct with homology arms upstream and downstream of *csxB* sandwiching the *ermB* cassette (pHF009).

For *bclA* deletion, homology arms complementary to regions immediately upstream and downstream of *bclA* were amplified by PCR (RF2153/2154, RF2157/2158 respectively) and the *ermB* cassette was amplified from pHF007 (RF2155/2156). The fragments were inserted into PCR linearised pHF012 (RF311/312) via Gibson assembly to produce the final construct with homology arms upstream and downstream of *bclA* sandwiching the *ermB* cassette (pHF015).

For the remaining constructs, *csxA*, *csxC* and *bclB* knockouts, inserts containing homology arms sandwiching the *ermB* cassette were synthesised by Genewiz (Azenta Life Sciences) and subsequently cloned between the BamHI and SacI sites in pJAK188 (*csxA*) or pHF012 (*csxC* and *bclB*) to produce constructs pHF010 (*csxA*), pHF013 (*csxC*), and pHF014 (*bclB*). All plasmid sequences were confirmed by Sanger sequencing.

To carry out mutagenesis, plasmids were transformed into *E. coli* CA434 and transferred to *C. sporogenes* by conjugation (Purdy 2002). Following conjugation, colonies were screened for single recombination via PCR and positive colonies were streaked to a lawn on non-selective BHI and incubated anaerobically for 2-3 days. Growth was harvested using 900 µl TY broth, used to inoculate 10 ml TY containing 4% (w/v) xylose and supplemented with erythromycin. This was incubated anaerobically for 8 h before 100 µl of a 10^−5^ and 10^−6^ dilution of the culture were streaked onto BHI containing 4% (w/v) xylose and erythromycin. Once growth occurred, 8-12 colonies were re-streaked to purity and screened for secondary recombination events via PCR. Strains were also screened for plasmid loss on selective plates containing thiamphenicol. Successful genomic mutations were further confirmed via Sanger sequencing.

### Electron Microscopy

#### Negative Staining

Spore samples (5 µl) were applied to glow discharged carbon coated copper grids. After 1 min the grid was blotted, washed once with distilled water, blotted, washed once with 0.75% (w/v) uranyl formate and then stained for 20 sec in 0.75% (w/v) uranyl formate. The grids were blotted and dried using a vacuum pump before imaging on a Philips CM100 transmission electron microscope (TEM), operated at 100kV, at magnifications between 950 - 3,900X equipped with a Gatan Multiscan 794 CCD camera.

#### Chemical Fixation

Spore pellets were fixed in 3% (w/v) glutaraldehyde (prepared in 0.1 M sodium cacodylate buffer) at 4 °C overnight. Samples were washed in 0.1 M sodium cacodylate and incubated for 2 hrs at room temperature in 1% (w/v) OsO_4_. Pellets were washed twice with 0.1 M sodium cacodylate and dehydrated by incubating with increasing concentrations of ethanol (50% (v/v), 75% (v/v), 95% (v/v), 100% (v/v) dried ethanol), followed by two pure propylene oxide 15 min incubations for complete dehydration. Samples were incubated overnight at room temperature in a 1:1 mix of 100% (v/v) propylene oxide and Epon resin to allow for infiltration. Resin was removed and excess propylene oxide evaporated at room temperature. Samples were incubated for two consecutive 4 hr periods in pure Epon resin before being embedded into the final fresh resin. Resin was polymerized by incubation at 60 °C for 48 hrs.

#### High Pressure Freezing and Freeze Substitution

High pressure freezing of *C. sporogenes* spores was carried out on a Leica EM PACT (Leica). Spore pellets were loaded into carriers and frozen in liquid nitrogen at pressures >2000 bar. The samples were then transferred into vials of freeze substitution cocktail (1% (w/v) OsO_4_, 0.5% (w/v) uranyl acetate, 3% (w/v) glutaraldehyde) at −90 °C. Freeze substitution (FS) was performed over 42 hrs in a Leica AFS2 using the following program: 12 hrs at −90 °C, 8 hrs gradual rise to −60 °C, 5 hrs at −60 °C, 7 hrs gradual rise to −30 °C, 5 hrs at −30 °C, and 5 hrs gradual rise to 0 °C. Freeze substituted samples were removed from the carriers by washing in acetone and treated with 1% uranyl acetate in acetone for 1 hr in the dark. Samples were washed 3 times for 30 min in 100% ethanol, followed by a final wash in acetone. Samples were then incubated in increasing concentrations of Spurr resin in acetone (4 hrs in 30% Spurr resin, overnight in 60%, 4 hrs in 90%, 2 x 2 hrs in 100%). Samples in 100% Spurr resin were placed into moulds and polymerised at 60 °C for 16 hrs.

#### Thin Sectioning

Resin thin sections (80-85 nm or 200-300 nm thickness) of spore samples were produced using an Ultracut E Ultramicrotome (Reichert-Jung). Sections were floated onto coated 300-square mesh nickel TEM grids (chemical fixation) or formvar coated 300-square mesh nickel TEM grids (freeze substitution). Sections were post-stained in 3% (w/v) uranyl acetate for 30 min, washed with dH_2_O, stained with Reynold’s lead citrate for 5 min and further washed with dH_2_O.

#### Imaging of 80-85 nm Sections

Sections were imaged on a Tecnai G2 spirit BioTwin (FEI) microscope, operated at 80 kV, at magnifications between 4,800 - 23,000X, equipped with a Gatan Orius SC1000B CCD camera.

#### Tomography of 200-300 nm Freeze Substituted Thin Sections

Tilt series were collected on a FEI Tecnai Arctica, equipped with a Falcon III direct electron detector (Thermo Fisher Scientific), operated at 200 kV. Grids were loaded at RT and cooled *in situ* with the autoloader to −195 °C. Tilt series were collected at a magnification of 16,600x (0.663 nm pixel size) in 3° increments between −60° and +60°, with a total dose of 60 e^−^/Å^2^.

#### Tomography Processing

Tilt series were aligned and reconstructed at a binning factor 4 using AreTomo3 v.2.2.2 (Zheng et al., 2022). 3dmod (IMOD 4.11.18)(Mastronarde and Held, 2017) was used for tomogram visualisation and analysis.

#### CsxA crystal preparation and imaging

Crystals of recombinant CsxA were grown as described by Janganan *et al*. (Janganan et al., 2020). 3 μl droplets of crystal suspension were applied to R 2/2 grids with a graphene oxide support (EM Resolutions), blotted for 5 s in a Leica EM GP plunge freezer and flash frozen in liquid ethane.

Images were recorded on a FEI Tecnai Arctica using the EPU software. Movie stacks were recorded in integration mode at 200 kV, 78,000X magnification with a defocus range of 1 - 6.5 μm. Specimen tilt angles were set to a nominal 0°, 20°, 40° and 60°.

#### CsxA crystal image processing

Movie frames were processed in MotionCor2 (Zheng et al., 2017). Images were processed within *2dx* (Gipson et al., 2007b; Gipson et al., 2007a) based on the MRC suite of programs (Henderson et al., 1986; Henderson et al., 1990; Crowther et al., 1996). Image defocus and crystal tilt angles were estimated with CTFFIND3 and CTFTILT (Mindell and Grigorieff, 2003) and lattice distortions corrected by unbending. Final structure factor phases and amplitudes were corrected for the effects of the contrast transfer function.

Plane group symmetry was determined from images of nominally untilted crystals in ALLSPACE (Valpuesta et al., 1994). A common phase origin was determined in ORIGTILTK, initially by comparing phases for 0° tilt images, followed by 20° images, then 40° images and finally 60° images (Amos et al., 1982). Initial phases and amplitudes along z* were estimated with LATLINE (Agard, 1983; Deatherage 1983). Cycles of crystal tilt angle and phase origin refinement were then performed against the LATLINE output in ORIGTILTK.

SCALIMAMP3D was used to correct image amplitudes before final merging (Havelka et al., 1995). Amplitudes and phases along the 0,0,l lattice line were approximated from a plot of contrast variation in Z of the initial calculated density map (Amos et al., 1982). Estimated 0,0,l amplitude values were scaled against the 1,0,0 values and a final density map calculated within the CCP4 software suite (Agirre et al., 2023).

#### Purification of surface layers from spores

0.2 g of spores were resuspended in 10 ml 0.02 M potassium phosphate buffer pH 7.2 and sonicated in 30 sec bursts for 40 mins total. The sample was spun at 2800 g at 4°C for 10 min and the supernatant containing spore surface layers was harvested. The remaining pellet was resuspended in a further 10 ml 0.02 M potassium phosphate buffer pH 7.2 and centrifuged at 1200 g at 4°C for 20 min. The supernatant was harvested and pooled with that of the previous spin. The spore surface material was then pelleted by centrifugation at 81,800 g at 4°C for 90 min and resusupended in 500 µl 0.02 M potassium phosphate buffer pH 7.2 before being added to the top of a 10-50% CsCl gradient. The sample was centrifuged at 50,000 g at 4°C for 16 hrs. Bands of material were isolated from the gradient and pelleted by centrifugation at 81,800 x g at 4°C for 90 mins. The pellets were washed a further 3 times by resuspension in 20 ml potassium phosphate buffer pH 7.5 and spun at 81,800 g at 4°C for 90 mins. Final pellets were resuspended in 100 µl TBS with 0.01% Triton and 3 µl were taken and screened by negative stain EM for crystal content.

#### Mass spectrometry analysis of spore surface layers

Samples from purified WT and Δ*csxA* spore surface layers were trypsin digested after cysteine derivatisation and subjected to nano-LC-MS/MS analysis. Proteins were identified using the proteome from the sequenced genome of *C. sporogenes* NCIMB 701792.

#### Data collection and image processing of surface layers from ΔcsxA spores

Purified spore surface layers from Δ*csxA* were applied to Quantifoil R 0.6/1 200 mesh gold grids and flash frozen on a Leica EM GP plunge freezer. Images were recorded on a 200 kV FEI Tecnai Arctica at 78,000X nominal magnification. We used the EPU software in “Lacey” mode to select areas for imaging prior to automated data collection. Underfocus was set in a range 1.5 to 3.5 μm.

Since parasporal layer crystals varied in thickness and tended to be rather mosaic, they were unsuitable for a standard 3D electron crystallography approach to image processing. Instead, we treated crystal patches as single particles and processed images within cryoSPARC (Punjani et al., 2017). After motion and CTF correction (see above), we used “blob picker” with minimum and maximum particle diameters of 55 and 75 Å respectively, for an initial set of particle picks. We used “inspect picks” to remove picks with very low or high local power, thus ensuring that most picks were within clearly crystalline areas. Particles were extracted in 256 X 256 boxes (pixel size 1.2 Å) and subjected to 2D class averaging. Class averages displaying a clear crystalline lattice were used for a second round of template-based particle picking and extraction with a 512 X 512 box. Subsequent 2D class averaging yielded one class with a clear crystalline lattice.

### Atomic Force Microscopy

Whole spores were adhered on poly-D-lysine coated slides (BD BioCoat poly-D-lysine coated 12 mm round tissue culture coverslips) or two different types of glass bottom petri dish: plain (FluoroDish, glass bottom, clear wall, 35 mm, 23 mm well, FD35-100) or custom made 50 μm grid glass bottom dishes (IBIDI, μ-Dish 35mm, low Grid-50 Glass Bottom, Ø 35 mm, low wall (800 μl volume), grid repeat distance 50 μm, #1.5 H (170 μm +/− 5 μm) D 263 M Schott glass) through incubation at an appropriate density in water for 30 min and rinsed with 5 x 1 ml of water. Both glass bottom petri dishes were pretreated with 0.01% poly-L-Lysine (Sigma-Aldrich). Purified spore surface layers from Δ*csxA* were adhered on poly-L-lysine (0.01%) treated cleaved mica (Agar Scientific) for 30 min and rinsed with 10 x 1 ml of water. Samples to be imaged in air were dried with a stream of filtered nitrogen. Samples to be imaged in water were used directly, in cases where an alternate buffer was used, buffer was exchanged following rinses with water. Samples that were dried and rehydrated were incubated in 3 ml of water for 30 mins before measurements.

Dried spores imaged in air were imaged with AC mode using a JPK Nanowizard 3 Ultra AFM, using NuNano Scout 350 RAI AFM probes (nominal spring constant and resonant frequency 42 N/m and 350 kHz, respectively) with a free amplitude of 20 nm and a setpoint of 65%, or a JPK Nanowizard 3 Bioscience AFM, with a free amplitude of 40 nm and a setpoint of approximately 70%. Images of the CsxC lattice on the spore surface were imaged with a free amplitude of 20 nm and a setpoint of approximately 65% on a JPK Nanowizard 3 Ultra. Height and phase images were obtained simultaneously.

Hydrated spores in water were imaged with Quantitative Imaging (QI) mode on a JPK Nanowizard 3 Bioscience AFM with a z length of 1 μm and a trigger force of 1 nN and BL-AC40TS AFM probes (Olympus) with a nominal spring constant and resonant frequency (in air) of 0.09 N/m and 110 kHz. Height images were collected at 80% of the trigger force.

Purified spore surface layers from Δ*csxA* were imaged with TappingMode™, in liquid, on a Bruker Dimension FastScan AFM with FastscanD probes (Bruker) with a nominal spring constant and resonant frequency of 0.25 N/m and 110 kHz (in liquid). The free amplitude was between 2-3 nm and a setpoint of approximately 90%.

Images were processed (cropping, flattening, plane fitting) using Gwyddion (Nečas and Klapetek, 2012).

### Calculation of volume from AFM images of dormant spores

Height images of individual spores were processed with flattening by masking the area of the spore, removing the polynomial background and aligning rows with median of differences in Gwyddion. The height scale was set to the full data range for each individual image and the image exported as an 8 bit tif. Depth and xy dimensions were recorded before export. These images were then processed in FIJI (Schindelin et al., 2012) with FIJI macro AFM Slicer (https://github.com/Laia-Pasquina/FIJI-AFMSlicer) using the surface and total apparent volume functions. Data and graphs were compiled in Origin(Pro), Version 2023b. OriginLab Corporation, Northampton, MA, USA.

## Abbreviations

A: appendage
AC: alternating current
AF: amorphous fringe
AFM: atomic force microscopy
BHI: brain heart infusion
C: core
CB: cuboid
CFU: colony forming units
CT: coat
CTF: contrast transfer function
CX: cortex
D: discontinuities
EM: electron microscopy
EX: exosporium
FE: folded edge
FS: freeze substitution
HN: hairy nap
IS: interspace
LB: Luria-Bertani broth
LP: lipped terminal protrusion
OD: optical density
PCR: polymerase chain reaction
PD: point defects
PL: parasporal layer
QI: quantitative imaging
RE: rough edge
RT: room temperature
SB: spore body
SD: screw dislocation
TEM: transmission electron microscopy
TY: tryptone yeast
WT: wild type

## Acknowledgements

This work was supported by BBSRC grant BB/W015072/1 to PAB, RF and JH and Wellcome Trust grant 221360/Z/20/Z to JH and NM. EM work was carried out in the University of Sheffield, School of Biosciences electron microscopy facility. The Dimension FastScan Bio AFM used in this work was purchased on BBSRC grant BB/L014904/1 to JH. We thank Chris Hill and Svetomir Tzokov for help with electron microscopy, Anamika Kumari and Theodora Enache for help with sample freezing for cryoEM, and Khoa Pham of the Faculty of Science Biological Mass Spectrometry Facility for assistance with MS analysis. Protein structure predictions were generated using AlphaFold3 (Google DeepMind). For manuscript preparation, ChatGPT (OpenAI, GPT-5.6 Sol,) was used to assist editing in the introduction and discussion and assist with literature searching. All structural models, text edits, and literature citations, were reviewed by the authors prior to submission.

## Author contributions

**Conceptualization:** Per Bullough, Robert Fagan, Jamie Hobbs

**Data curation:** Per Bullough, Robert Fagan, Jamie Hobbs

**Formal analysis:** Per Bullough, Hannah Fisher, Abigail Roberts, Nic Mullin

**Funding acquisition:** Per Bullough, Robert Fagan, Jamie Hobbs

**Investigation:** Hannah Fisher, Abigail Roberts, Ainhoa Dafis-Sagarmendi, Nic Mullin, Jessica Buddle, Thamarai Janganan, Per Bullough

**Methodology:** Hannah Fisher, Abigail Roberts, Ainhoa Dafis-Sagarmendi, Nic Mullin, Jessica Buddle, Thamarai Janganan, Per Bullough

**Project administration:** Per Bullough, Robert Fagan, Jamie Hobbs

**Resources:** Robert Fagan

**Supervision:** Per Bullough, Robert Fagan, Jamie Hobbs

**Validation:** Per Bullough, Robert Fagan, Jamie Hobbs

**Visualization:** Hannah Fisher, Abigail Roberts, Per Bullough

**Writing – original draft:** Hannah Fisher, Abigail Roberts, Jessica Buddle, Per Bullough

**Writing – review & editing:** Hannah Fisher, Abigail Roberts, Ainhoa Dafis-Sagarmendi, Nic Mullin, Jessica Buddle, Thamarai Janganan, Jamie Hobbs, Robert Fagan, Per Bullough

## Supporting information

**S1 Fig.**
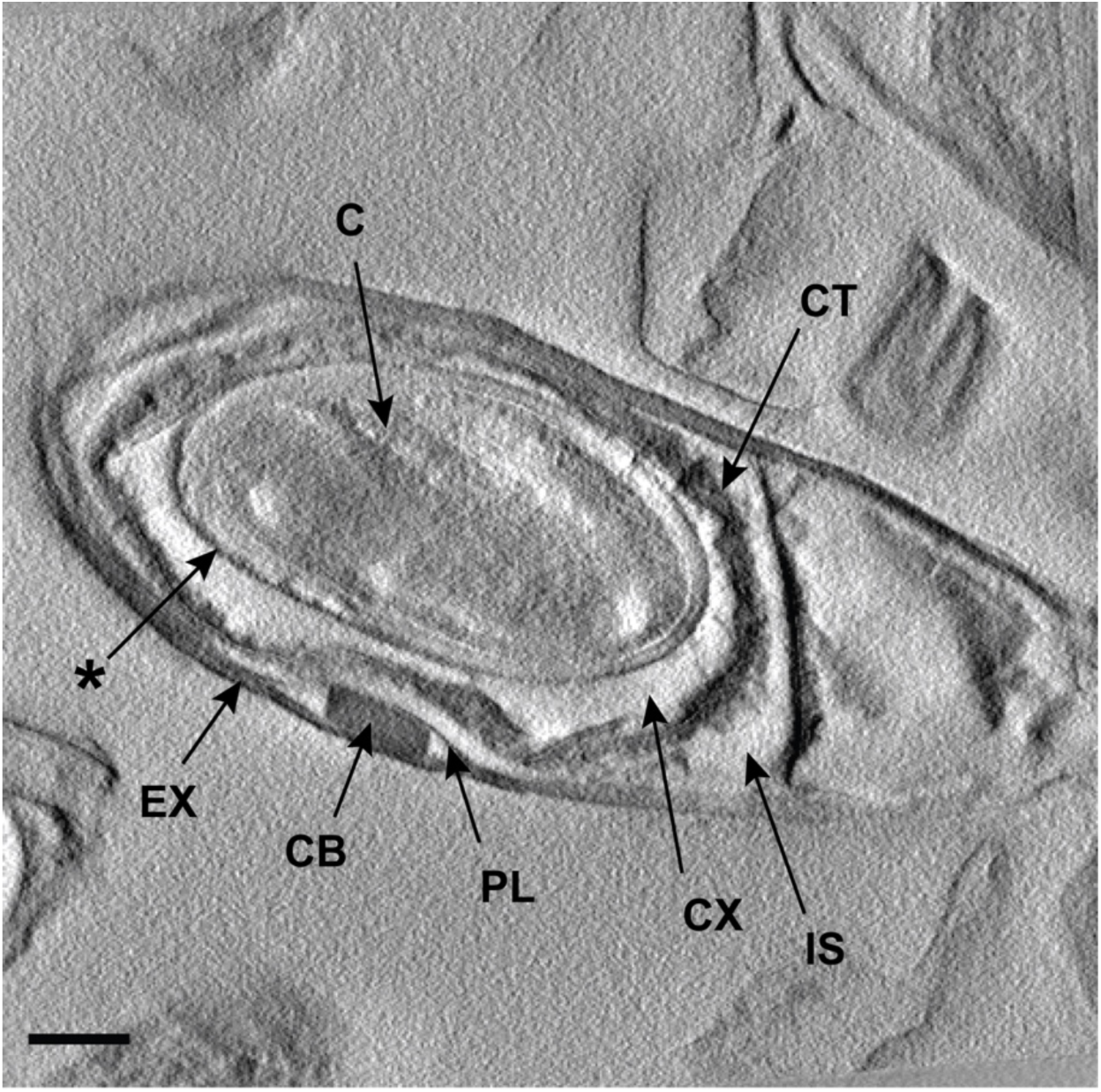
Electron tomography provides a three-dimensional view of the spore interior. Projection calculated from a 13 nm slice of a tomogram from a freeze-substituted spore. The full tomogram can be seen in S1 Movie. C - core, CX - cortex, CT - coat, IS - interspace, PL - parasporal layers, CB - cuboid, EX - exosporium, **\*** - unidentified layer

**S2 Fig.**
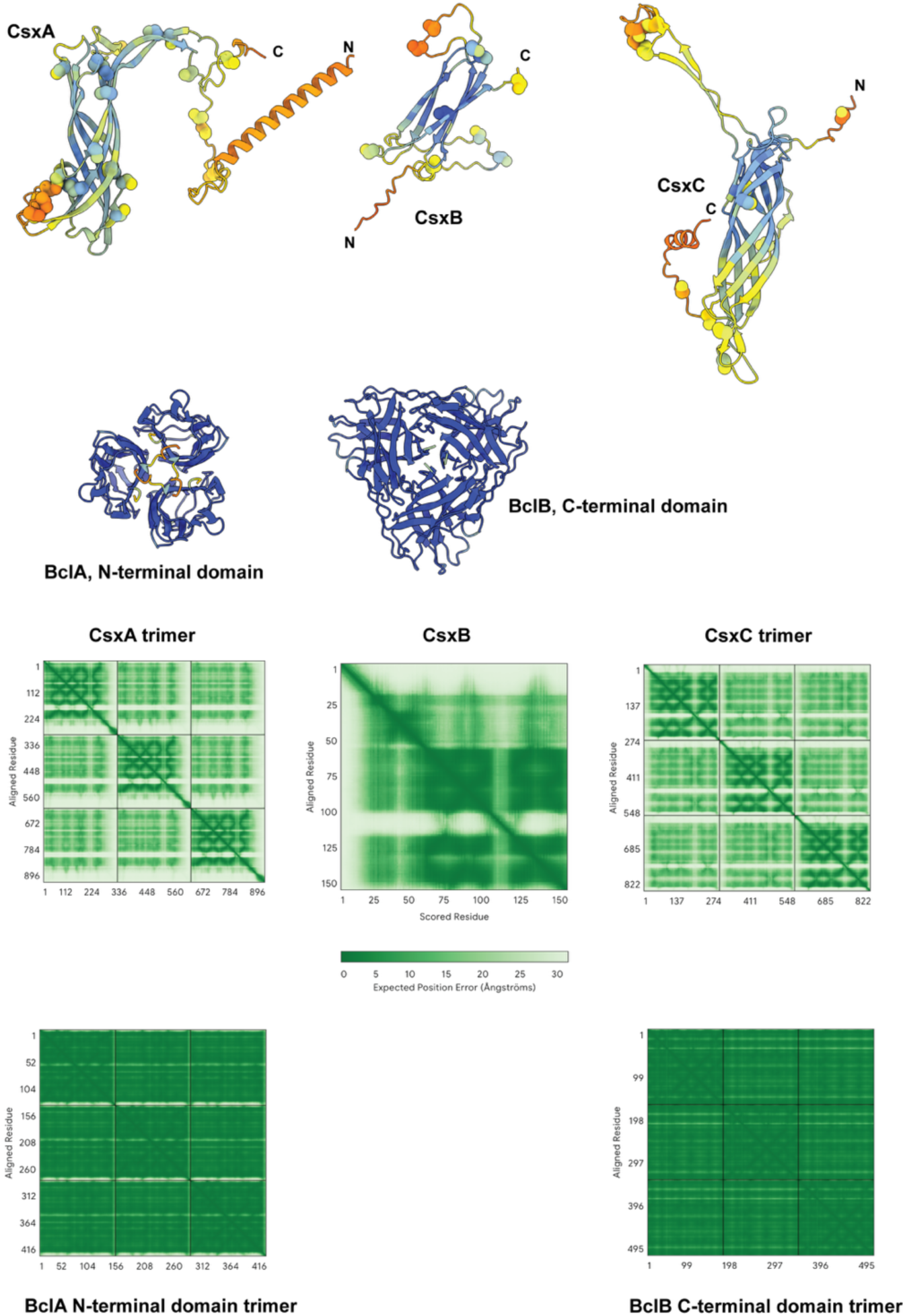
AlphaFold3 predictions of *C. sporogenes* spore envelope proteins. For CsxA and CsxC, only single subunits from predicted trimers are shown. Full predicted timers are shown in Fig 5. Cysteine residues are shown as large spheres. Predicted trimers are shown for BclA and BclB domains. plDDT scores are shown in colour from ‘very low confidence’ in amber to ‘very high confidence’ in dark blue. Expected positional errors are shown for trimeric models with full sequence for CsxA and CsxC, for the full monomer sequence of CsxB and respective terminal domain sequences only for BclA and BclB.

**S3 Fig.**
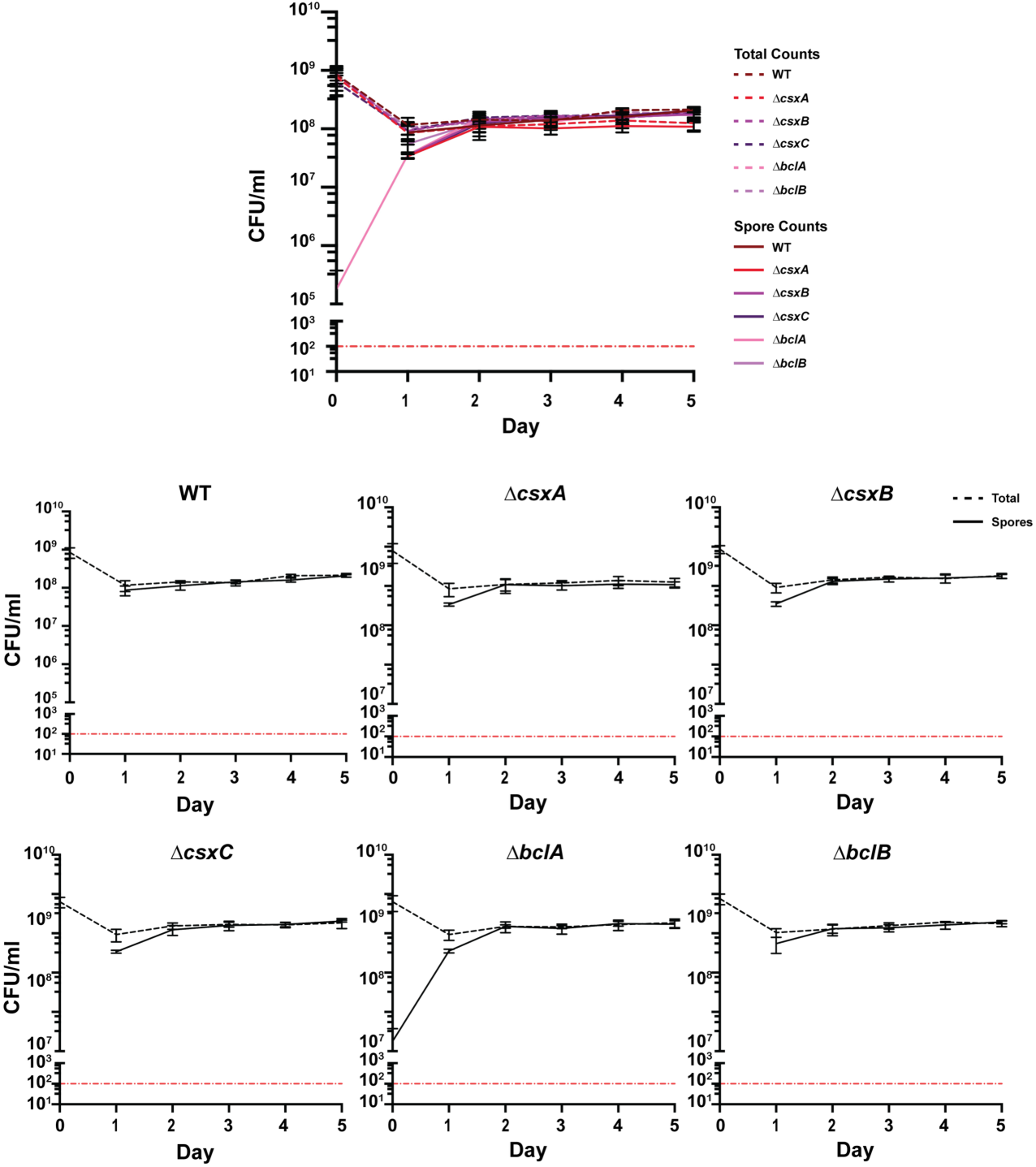
Sporulation assay of WT and exosporium mutant strains. Total cell counts were determined by enumeration of CFUs on TY agar. Spore counts were determined likewise following a 30 min incubation period at 65 °C. Experiments were conducted in triplicate on biological duplicates. The means ± standard deviation (error bars) are shown.

**S4 Fig.**
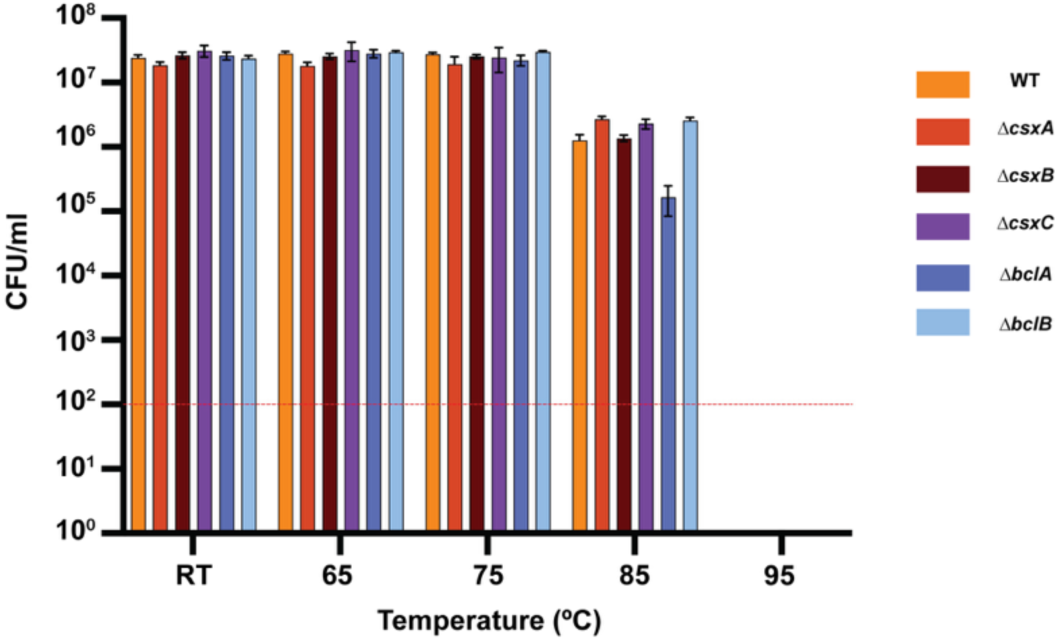
Spore viability after heat treatment of WT and exosporium mutant spores. CFUs were enumerated following incubation at each temperature for 30 min. Experiments were conducted in triplicate on biological duplicates. The means ± standard deviations (error bars) are shown.

**S5 Fig.**
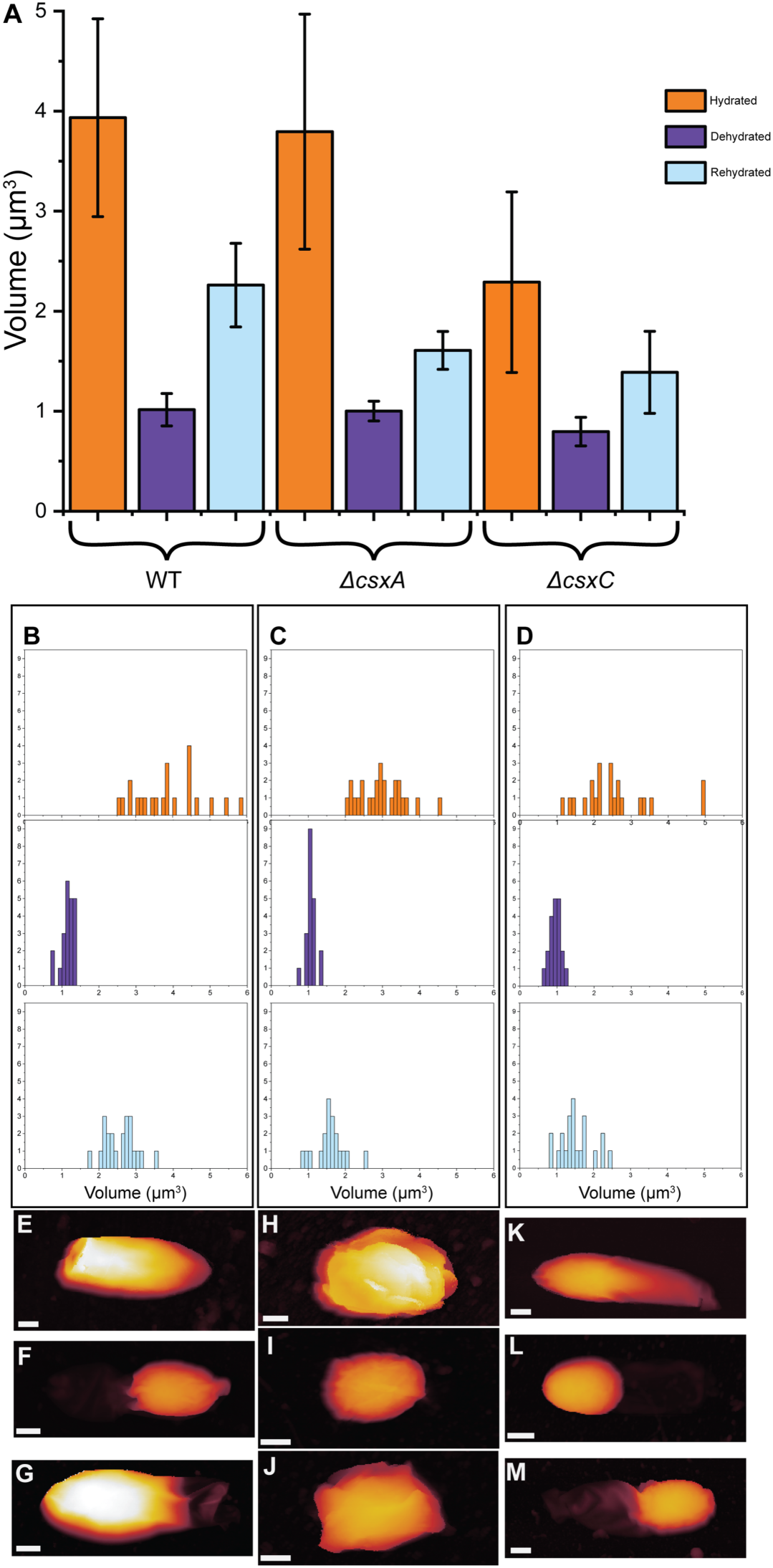
Changes in spore volume and appearance during hydration cycling. **A** Average volume calculated from AFM height images of individual spores using Fiji macro AFM Slicer (https://github.com/Laia-Pasquina/FIJI-AFMSlicer) which calculates the enclosed volume from sequential height slices through the reconstructed topography, N ≥ 19. Scale bars represent standard deviation. **B-D** The distribution of individual spore volume measurements of hydrated (orange), dehydrated (purple), and rehydrated (blue) for WT (**B**), Δ*csxA* (**C**), and Δ*csxC* **(D)**. **E - M** Example height images of spores in a hydrated **(E, H, K)**, dehydrated **(F, I, L)**, and rehydrated **(G, J, M)** for WT **(E-G)**, Δ*csxA* **(H-J)**, and Δ*csxC* **(K-M)** spores. Height range: 1.25 μm. Scale bar: 500 nm.

**S6 Fig.**
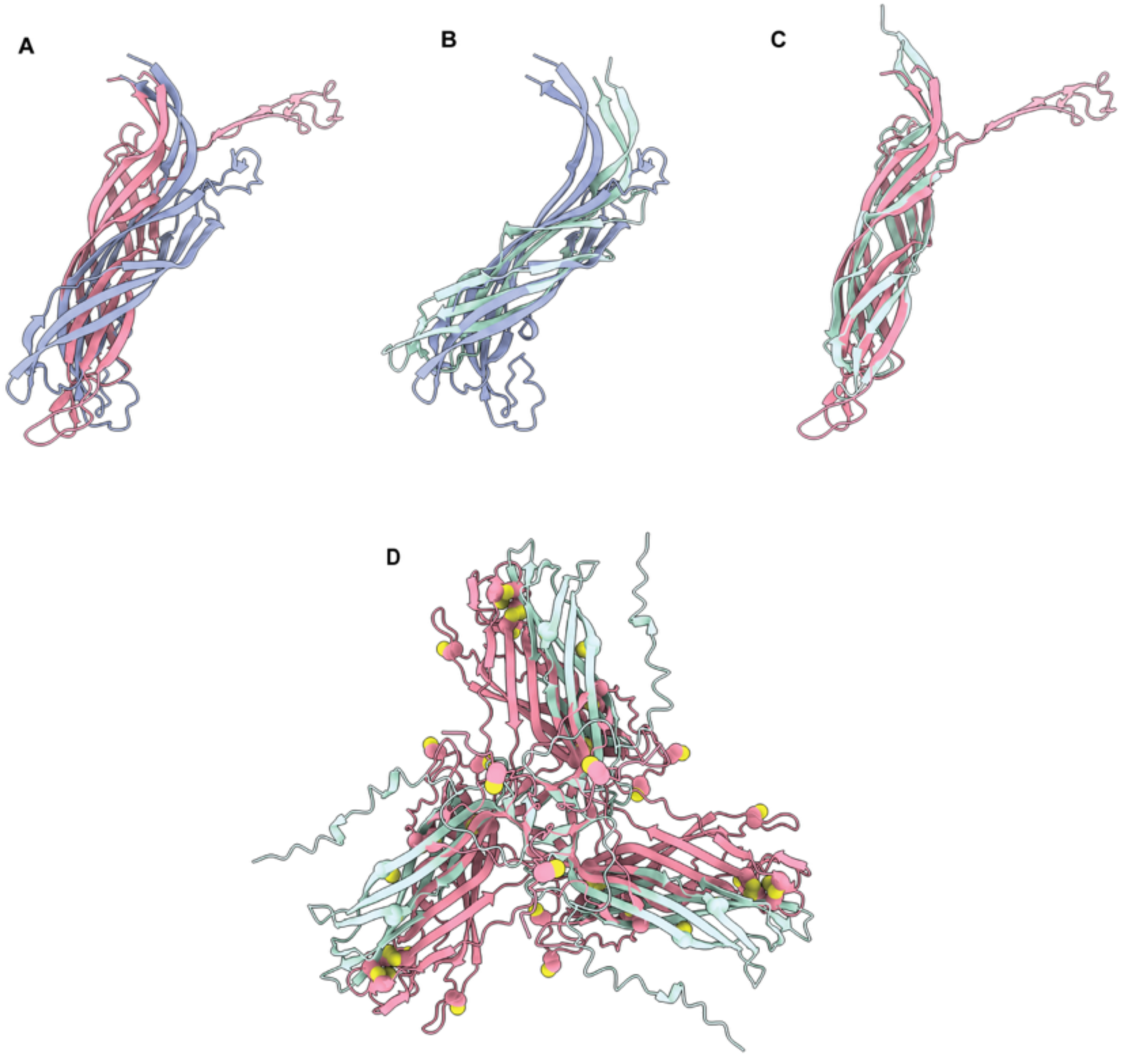
Predicted CsxA and CsxC DUF3794/SPOCS domains align with CotE. **A** Alignment of CsxA (mauve) and CsxC (salmon) single subunits. **B** Alignment of CsxA (mauve) and CotE (pale blue). **C** Alignment of CsxC (salmon) and CotE (pale blue). In (A)-(C), unstructured N- and C-terminal regions have been deleted. **D** Alignment of CsxC (salmon) and CotE (pale blue) on all residues of the predicted trimer. Cysteine residues are shown as spheres, with the vast majority being in CsxC.

**S7 Fig.**
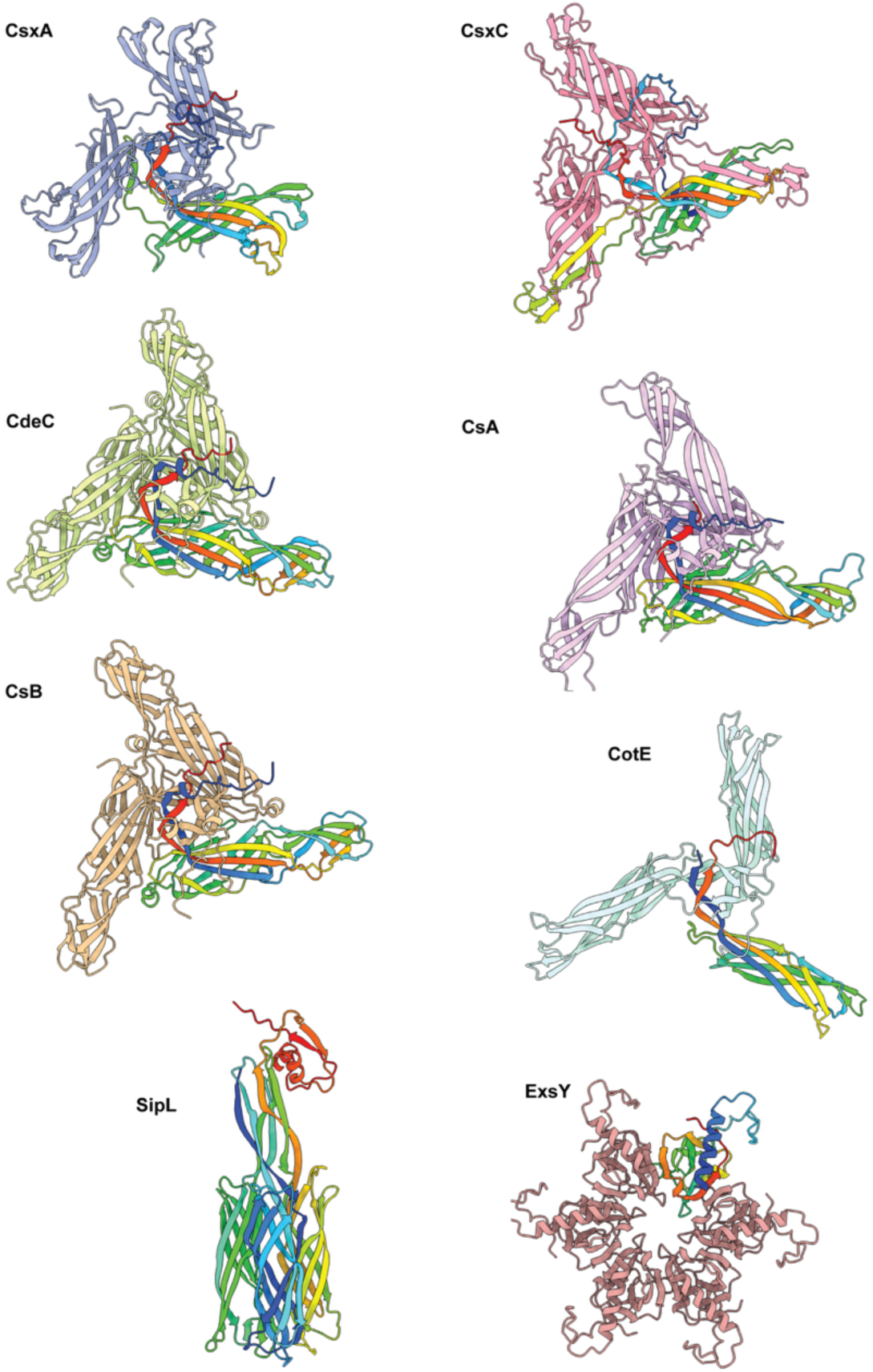
Alphafold3 predictions of spore proteins from other species reveal similarities within Clostridia and differences from Bacilli. *C. sporogenes* CsxA and CsxC compared with *C. difficile* CdeC, *C. sordellii* CsA and CsB, *B. subtilis* CotE and SipL and *B. cereus* ExsY. For clarity, unstructured N- and C-terminal residues have been removed. In each multimer, one subunit has been coloured from remaining N-terminal residues (blue) to C-terminal residues (red).

**S1 Movie. Tomogram of dormant WT spore.** Full tomogram reconstruction from a 85 mm freeze-substituted then section of a dormant WT spore provides a three-dimensional view of interior spore features. Scale bar 250 nm.

https://drive.google.com/file/d/1O8xjPBFR4zp_xC3O5n0ADH1Pe1muHmSd/view?usp=drive_link

**S2 Movie. CsxA reconstruction and modelling.** A fit of the CsxA lattice model into the cryoEM reconstruction.

https://drive.google.com/file/d/1rW8D2w2R8dqznWHuml4caQsarzLy0_Mx/view?usp=drive_link

**S1 Table. Confidence of predicted disulphide pairs in a trimeric CsxA assembly.**

https://docs.google.com/document/d/1SCqPGiWIqJvajH5dXAZHhzsRyek9y_PM/edit?usp=drive_link&ouid=109212407047640205494&rtpof=true&sd=true

**S2 Table. Confidence of predicted disulphide pairs in a trimeric CsxC assembly.**

https://docs.google.com/document/d/1zU3-pz0yGKbVxN6ZfaxrTA6rE0cRfHOo/edit?usp=drive_link&ouid=109212407047640205494&rtpof=true&sd=true

**S3 Table. Data collection and LATLINE processing statistics for CryoEM of CsxA 2D crystals.**

https://docs.google.com/document/d/1QAMhrhkwejTmSU7XDiGEBGiz_YAMDqra/edit?usp=drive_link&ouid=109212407047640205494&rtpof=true&sd=true

**S4 Table. Strains used in this study.**

https://docs.google.com/document/d/1tDAKntj5w7tsrfA28UIaAXiyeU7aHeZ6/edit?usp=drive_link&ouid=109212407047640205494&rtpof=true&sd=true

**S5 Table. Plasmids used in this study.**

https://docs.google.com/document/d/1Ja4yZkMAsFDRxTjdgy7h-f9H81lAqBm2/edit?usp=drive_link&ouid=109212407047640205494&rtpof=true&sd=true

**S6 Table 6. Primers used in this study**

https://docs.google.com/document/d/1K07CqH9LomwdLCJP1VknVz8ezC1ks9Jv/edit?usp=drive_link&ouid=109212407047640205494&rtpof=true&sd=true

**S1 Data. Mass spectrometry data**

https://docs.google.com/spreadsheets/d/1-wk9xp4Lsy6WS_38md-GZRfxdRVeArN3/edit?usp=drive_link&ouid=109212407047640205494&rtpof=true&sd=true

## References

Abramson, J., Adler, J., Dunger, J., Evans, R., Green, T., Pritzel, A., et al. (2024) Accurate structure prediction of biomolecular interactions with AlphaFold 3. Nature 630: 493–500.

Agard, D.A. (1983) A Least-Squares Method for Determining Structure Factors in 3-Dimensional Tilted-View Reconstructions. Journal of Molecular Biology 167: 849–852.

Agirre, J., Atanasova, M., Bagdonas, H., Ballard, C.B., Baslé, A., Beilsten-Edmands, J., et al. (2023) The CCP4 suite: integrative software for macromolecular crystallography. Acta Crystallogr Sect D 79: 449–461.

Amos, L.A., Henderson, R., and Unwin, P.N.T. (1982) Three-dimensional structure determination by electron microscopy of two-dimensional crystals. Prog Biophys Mol Biol 39: 183–231.

Antunes, W., Pereira, F.C., Feliciano, C., Saujet, L., Vultos, T. dos, Couture-Tosi, E., et al. (2018) Structure and assembly of a *Clostridioides difficile* spore polar appendage. bioRxiv 468637.

Arnon, S.S. (1977) Infant Botulism: Epidemiological, Clinical, and Laboratory Aspects. JAMA 237: 1946.

Aronson, A.I., Ekanayake, L., and Fitz-James, P.C. (1992) Protein filaments may initiate the assembly of the *Bacillus subtilis* spore coat. Biochimie 74: 661–667

Ball, D.A., Taylor, R., Todd, S.J., Redmond, C., Couture-Tosi, E., Sylvestre, P., et al. (2008) Structure of the exosporium and sublayers of spores of the *Bacillus cereus* family revealed by electron crystallography. Mol Microbiol 68: 947–958

Bankevich, A., Nurk, S., Antipov, D., Gurevich, A.A., Dvorkin, M., Kulikov, A.S., et al. (2012) SPAdes: A New Genome Assembly Algorithm and Its Applications to Single-Cell Sequencing. J Comput Biol 19: 455–477.

Barra-Carrasco, J., Olguin-Araneda, V., Plaza-Garrido, A., Miranda-Cardenas, C., Cofre-Araneda, G., Pizarro-Guajardo, M., et al. (2013) The *Clostridium difficile* Exosporium Cysteine (CdeC)-Rich Protein Is Required for Exosporium Morphogenesis and Coat Assembly. J Bacteriol 195: 3863–3875.

Barwinska-Sendra, A., Salgado, P.S., and Sendra, K.M. (2025) Evolutionary plasticity of bacterial surface layer protein exoskeletons. bioRxiv 2025.04.02.646754.

Bauda, E., Gallet, B., Moravcova, J., Effantin, G., Chan, H., Novacek, J., et al. (2024) Ultrastructure of macromolecular assemblies contributing to bacterial spore resistance revealed by in situ cryo-electron tomography. Nat Commun 15: 1376.

Bauer, T., Little, S., Stöver, A.G., and Driks, A. (1999) Functional regions of the *Bacillus subtilis* spore coat morphogenetic protein CotE. Journal of Bacteriology 181: 7043–7051

Bolger, A.M., Lohse, M., and Usadel, B. (2014) Trimmomatic: a flexible trimmer for Illumina sequence data. Bioinformatics 30: 2114–2120.

Boone, T.J., Mallozzi, M., Nelson, A., Thompson, B., Khemmani, M., Lehmann, D., et al. (2018) Coordinated Assembly of the *Bacillus anthracis* Coat and Exosporium during Bacterial Spore Outer Layer Formation. mBio 9: e01166–18.

Brunt, J., Cross, K.L., and Peck, M.W. (2015) Apertures in the *Clostridium sporogenes* spore coat and exosporium align to facilitate emergence of the vegetative cell. Food Microbiol 51: 45–50.

Cartman, S.T, Kelly, M.L., Heeg, D., and Minto, N.P. (2012) Precise Manipulation of the *Clostridium difficile* Chromosome Reveals a Lack of Association between the tcdC Genotype and Toxin Production. Appl Env Biol 78: 4683–4690.

Crowther, R.A., Henderson, R., and Smith, J.M. (1996) MRC Image Processing Programs. J Struct Biol 116: 9–16.

Davidson, P., Eutsey, R., Redler, B., Hiller, N.L., Laub, M.T., and Durand, D. (2018) Flexibility and constraint: Evolutionary remodeling of the sporulation initiation pathway in Firmicutes. PLoS Genet 14: e1007470.

Deatherage, J.F., Taylor, K.A., and Amos, L.A. (1983) Three-dimensional arrangement of the cell wall protein of *Sulfolubus acidocaldarius*. J Mol Biol 167: 823–852.

Delerue, T., Anantharaman, V., Gilmore, M.C., Popham, D.L., Cava, F., Aravind, L., and Ramamurthi, K.S. (2022) Bacterial developmental checkpoint that directly monitors cell surface morphogenesis. Dev Cell 57: 344–360.e6.

Dembek, M., Barquist, L., Boinett, C.J., Cain, A.K., Mayho, M., Lawley, T.D., et al. (2015) High-Throughput Analysis of Gene Essentiality and Sporulation in *Clostridium difficile*. mBio 6: e02383–14.

Driks, A. (1999) *Bacillus subtilis* spore coat. Microbiology and Molecular Biology Reviews : MMBR 63: 1–20

Driks, A., and Eichenberger, P. (2016) The Spore Coat. Microbiol Spectr 4: 1–22.

Ebersold, H.R., Lüthy, P., Cordier, J.L., and Müller, M. (1981) A Freeze-Substitution and Freeze-Fracture Study of Bacterial Spore Structures. Journal of ultrastructure research 76: 71–81.

Fagan, R.P., and Fairweather, N.F. (2014) Biogenesis and functions of bacterial S-layers. Nat Rev Microbiol 12: 211–222.

Fisher, H., Kubisa, L. de L., Jakhmola, A., Walker, E., Kirk, J.A., Oatley, P., et al. (2026) A set of genetic tools for use in *Clostridioides difficile* and related species. Microbiology 172.

Frank, F.C. (1949) The influence of dislocations on crystal growth. Discuss Faraday Soc 5: 48–54.

Fuchs, M., Lamm-Schmidt, V., Sulzer, J., Ponath, F., Jenniches, L., Kirk, J.A., et al. (2021) An RNA-centric global view of *Clostridioides difficile* reveals broad activity of Hfq in a clinically important gram-positive bacterium. Proc Natl Acad Sci 118: e2103579118.

Galperin, M.Y., Mekhedov, S.L., Puigbo, P., Smirnov, S., Wolf, Y.I., and Rigden, D.J. (2012) Genomic determinants of sporulation in Bacilli and Clostridia: towards the minimal set of sporulation-specific genes. Environ Microbiol 14: 2870–2890.

Galperin, M.Y., Yutin, N., Wolf, Y.I., Alvarez, R.V., and Koonin, E.V. (2022) Conservation and Evolution of the Sporulation Gene Set in Diverse Members of the Firmicutes. J Bacteriol 204: e00079–22.

Gipson, B., Zeng, X., and Stahlberg, H. (2007a) 2dx_merge: Data management and merging for 2D crystal images. J Struct Biol 160: 375–384.

Gipson, B., Zeng, X., Zhang, Z.Y., and Stahlberg, H. (2007b) 2dx - User-friendly image processing for 2D crystals. J Struct Biol 157: 64–72.

Gould, G.W., Stubbs, J.M., and King, W.L. (1970) Structure and Composition of Resistant Layers in Bacterial Spore Coats. Microbiology 60: 347–355.

Gurevich, A., Saveliev, V., Vyahhi, N., and Tesler, G. (2013) QUAST: quality assessment tool for genome assemblies. Bioinformatics 29: 1072–1075.

Havelka, W.A., Henderson, R., and Oesterhelt, D. (1995) Three-dimensional structure of halorhodopsin at 7 Å resolution. J Mol Biol 247: 726–738.

Henderson, R., Baldwin, J., Downing, K., Lepault, J., and Zemlin, F. (1986) Structure of purple membrane from *Halobacterium halobium*: recording, measurement and evaluation of electron micrographs at 3.5 A resolution. Ultramicroscopy 19: 147–178.

Henderson, R., Baldwin, J.M., Ceska, T.A., Zemlin, F., Beckmann, E., and Downing, K.H. (1990) Model for the structure of bacteriorhodopsin based on high-resolution electron cryo-microscopy. J Mol Biol 213: 899–929.

Henriques, A.O., and Jr., C.P.M. (2007) Structure, Assembly, and Function of the Spore Surface Layers. Microbiology 61: 555–588

Henriques, A.O., and Moran, C.P. (2000) Structure and Assembly of the Bacterial Endospore Coat. Methods 20: 95–110

Hoeniger, J.F., and Headley, C.L. (1969) Ultrastructural aspects of spore germination and outgrowth in *Clostridium sporogenes*. Canadian journal of microbiology 15: 1061–1065.

Holser, W.T. (1958) Point Groups and Plane Groups in a Two-Sided Plane and their Subgroups. Z für Krist 110: 266–281.

Janganan, T.K., Mullin, N., Dafis-Sagarmendi, A., Brunt, J., Tzokov, S.B., Stringer, S., et al. (2020) Architecture and Self-Assembly of *Clostridium sporogenes* and *Clostridium botulinum* Spore Surfaces Illustrate a General Protective Strategy across Spore Formers. mSphere 5: e00424–20.

Janganan, T.K., Mullin, N., Tzokov, S.B., Stringer, S., Fagan, R.P., Hobbs, J.K., et al. (2016) Characterization of the spore surface and exosporium proteins of *Clostridium sporogenes*; implications for *Clostridium botulinum* group I strains. Food Microbiol 59: 205–212.

Jiang, S., Wan, Q., Krajcikova, D., Tang, J., Tzokov, S.B., Barak, I., and Bullough, P.A. (2015) Diverse supramolecular structures formed by self-assembling proteins of the *Bacillus subtilis* spore coat. Mol Microbiol 97: 347–359

Johnston, E., Isbilir, B., Alva, V., Bharat, T.A.M, and Doye, J.P.K. (2024) Punctuated and continuous structural diversity of S-layers across the prokaryotic tree of life. bioRxiv.

Kailas, L., Terry, C., Abbott, N., Taylor, R., Mullin, N., Tzokov, S.B., et al. (2011) Surface architecture of endospores of the *Bacillus cereus*/*anthracis*/*thuringiensis* family at the subnanometer scale. Proc Natl Acad Sci 108: 16014–16019.

Kempen, M. van, Kim, S.S., Tumescheit, C., Mirdita, M., Lee, J., Gilchrist, C.L.M., et al. (2024) Fast and accurate protein structure search with Foldseek. Nat Biotechnol 42: 243–246.

Kirchdoerfer, R.N., Herrin, B.R., Han, B.W., Turnbough, C.L., Cooper, M.D., and Wilson, I.A. (2012) Variable Lymphocyte Receptor Recognition of the Immunodominant Glycoprotein of *Bacillus anthracis* Spores. Structure 20: 479–486.

Koch, R. (1876). Die Ätiologie der Milzbrand-Krankheit, begründet auf die Entwicklungsgeschichte des *Bacillus anthracis*. Beiträge zur Biologie der Pflanzen, 2, 277–310.

Lee, D., Baek, Y., Park, M., Kim, D., Byun, K., Hyun, J., and Ha, N.-C. (2025) 3D meshwork architecture of the outer coat protein CotE: implications for bacterial endospore sporulation and germination. mBio 16: e02472–24.

Lehmann, D., Sladek, M., Khemmani, M., Boone, T.J., Rees, E., and Driks, A. (2022) Role of novel polysaccharide layers in assembly of the exosporium, the outermost protein layer of the *Bacillus anthracis* spore. Mol Microbiol 118: 258–277.

Little, S., and Driks, A. (2001) Functional analysis of the *Bacillus subtilis* morphogenetic spore coat protein CotE. Molecular Microbiology 42: 1107–1120.

Lund, B., Gee, J., King, N., Horne, R., and Harnden, J. (1978) The Structure of the Exosporium of a Pigmented Clostridium. Journal of general microbiology 105: 165–174.

Mackey, B.M., and Morris, J.G. (1972) The exosporium of *Clostridium pasteurianum*. Journal of general microbiology 73: 325–338

Mackey, B.M. and Morris, J.G. (1972) The Exosporium of *Clostridium pasteurianum*. J Gen Microbiol 73: 325–338.

Malkin, A.J., Kuznetsov, Yu.G., Land, T.A., DeYoreo, J.J., and McPherson, A. (1995) Mechanisms of growth for protein and virus crystals. Nat Struct Biol 2: 956–959.

Mastronarde, D.N., and Held, S.R. (2017) Automated tilt series alignment and tomographic reconstruction in IMOD. J Struct Biol 197: 102–113.

Masuda, K., Kawata, T., Takumi, K., and Kinouchi, T. (1980) Ultrastructure of a hexagonal array in exosporium of a highly sporogenic mutant of *Clostridium botulinum* type A revealed by electron microscopy using optical diffraction and filtration. Microbiology and immunology 24: 507–513.

McKenney, P.T., Driks, A., and Eichenberger, P. (2013) The *Bacillus subtilis* endospore: assembly and functions of the multilayered coat. Nat Rev Microbiol 11: 33–44.

Mindell, J.A., and Grigorieff, N. (2003) Accurate determination of local defocus and specimen tilt in electron microscopy. J Struct Biol 142: 334–47.

Mistry, J., Chuguransky, S., Williams, L., Qureshi, M., Salazar, G.A., Sonnhammer, E.L.L., et al. (2021) Pfam: The protein families database in 2021. Nucleic acids Res 49: D412–D419.

Nečas, D., and Klapetek, P. (2012) Gwyddion: an open-source software for SPM data analysis. Cent Eur J Phys 10: 181–188.

Oatley, P., Kirk, J.A., Ma, S., Jones, S., and Fagan, R.P. (2020) Spatial organization of *Clostridium difficile* S-layer biogenesis. Sci Rep 10: 14089.

Panessawarren, B.J., Tortora, G.T., and Warren, J.B. (1994) Electron-Microscopy of *Clostridium sporogenes* Endospore Attachment and Germination. Scanning 16: 227–240.

Paysan-Lafosse, T., Andreeva, A., Blum, M., Chuguransky, S.R., Grego, T., Pinto, B.L., et al. (2024) The Pfam protein families database: embracing AI/ML. Nucleic Acids Res 53: D523–D534.

Plomp, M., McCaffery, J.M., Cheong, I., Huang, X., Bettegowda, C., Kinzler, K.W., et al. (2007) Spore Coat Architecture of *Clostridium novyi* NT Spores. J Bacteriol 189: 6457–6468.

Portinha, I.M., Douillard, F.P., Korkeala, H., and Lindström, M. (2022) Sporulation Strategies and Potential Role of the Exosporium in Survival and Persistence of *Clostridium botulinum*. Int J Mol Sci 23: 754.

Pum, D., Breitwieser, A., and Sleytr, U.B. (2021) Patterns in Nature - S-Layer Lattices of Bacterial and Archaeal Cells. Crystals 11: 869.

Punjani, A., Rubinstein, J.L., Fleet, D.J., and Brubaker, M.A. (2017) cryoSPARC: algorithms for rapid unsupervised cryo-EM structure determination. Nat Methods 14: 290–296.

Putnam, E.E., Nock, A.M., Lawley, T.D., and Shen, A. (2013) SpoIVA and SipL are *Clostridium difficile* Spore Morphogenetic Proteins. J Bacteriol 195: 1214–1225.

Rabi, R., Larcombe, S., Mathias, R., McGowan, S., Awad, M., and Lyras, D. (2018) *Clostridium sordellii* outer spore proteins maintain spore structural integrity and promote bacterial clearance from the gastrointestinal tract. PLoS Pathog 14: e1007004.

Rabi, R., Turnbull, L., Whitchurch, C.B., Awad, M., and Lyras, D. (2017) Structural Characterization of *Clostridium sordellii* Spores of Diverse Human, Animal, and Environmental Origin and Comparison to *Clostridium difficile* Spores. mSphere 2: e00343–17.

Réty, S., Salamitou, S., Garcia-Verdugo, I., Hulmes, D.J.S., Hégarat, F.L., Chaby, R., and Lewit-Bentley, A. (2005) The Crystal Structure of the *Bacillus anthracis* Spore Surface Protein BclA Shows Remarkable Similarity to Mammalian Proteins. J Biol Chem 280: 43073–43078

Reviakine, I., Georgiou, D.K., and Vekilov, P.G. (2003) Capillarity Effects on Crystallization Kinetics: Insulin. J Am Chem Soc 125: 11684–11693.

Romero-Rodríguez, A., Troncoso-Cotal, S., Guerrero-Araya, E., and Paredes-Sabja, D. (2020) The *Clostridioides difficile* Cysteine-Rich Exosporium Morphogenetic Protein, CdeC, Exhibits Self-Assembly Properties That Lead to Organized Inclusion Bodies in *Escherichia coli*. mSphere 5: e01065–20.

Schindelin, J., Arganda-Carreras, I., Frise, E., Kaynig, V., Longair, M., Pietzsch, T., et al. (2012) Fiji: an open-source platform for biological-image analysis. Nat Methods 9: 676–682.

Secaira-Morocho, H., Castillo, J.A., and Driks, A. (2020) Diversity and evolutionary dynamics of spore-coat proteins in spore-forming species of Bacillales. Microb Genom 6: mgen000451.

Seemann, T. (2014) Prokka: rapid prokaryotic genome annotation. Bioinformatics 30: 2068–2069.

Setlow, P. (2014a) Germination of Spores of Bacillus Species: What We Know and Do Not Know. J Bacteriol 196: 1297–1305.

Setlow, P. (2014b) Spore Resistance Properties. Microbiol Spectr 2: TBS-0003-2012.

Sorg, J.A., and Sonenshein, A.L. (2008) Bile Salts and Glycine as Cogerminants for *Clostridium difficile* Spores. J Bacteriol 190: 2505–2512.

Stevenson, K.E., and Vaughn, R.H. (1972) Variability in ultrastructure of *Clostridium botulinum* spores. Can J Microbiol 18: 1717–1719.

Stewart, G.C. (2015) The Exosporium Layer of Bacterial Spores: a Connection to the Environment and the Infected Host. Microbiol Mol Biol Rev 79: 437–457.

Stewart, G.C. (2017) Assembly of the outermost spore layer: pieces of the puzzle are coming together. Mol Microbiol 104: 535–538.

Sylvestre, P., Couture-Tosi, E., and Mock, M. (2003) Polymorphism in the Collagen-Like Region of the *Bacillus anthracis* BclA Protein Leads to Variation in Exosporium Filament Length. J Bacteriol 185: 1555–1563.

Takumi, K., Kinouchi, T., and Kawata, T. (1979) Isolation and Partial Characterization of Exosporium from Spores of a Highly Sporogenic Mutant of *Clostridium botulinum* Type A. Microbiol Immunol 23: 443–454.

Terry, C., Jiang, S., Radford, D.S., Wan, Q., Tzokov, S., Moir, A., and Bullough, P.A. (2017) Molecular tiling on the surface of a bacterial spore – the exosporium of the *Bacillus anthracis*/*cereus*/*thuringiensis* group. Mol Microbiol 104: 539–552.

Touchette, M.H., Puebla, H.B. de la, Ravichandran, P., and Shen, A. (2019) SpoIVA-SipL Complex Formation Is Essential for *Clostridioides difficile* Spore Assembly. J Bacteriol 201: 10.1128/jb.00042-19.

Valpuesta, J.M., Carrascosa, J.L., and Henderson, R. (1994) Analysis of Electron Microscope Images and Electron Diffraction Patterns of Thin Crystals of Ø29 Connectors in Ice. J Mol Biol 240: 281–287.

Walker, P.D., Thomson, R.O., and Baillie, A. (1967) Fine Structure of Clostridia with Special Reference to the Location of Antigens and Enzymes. J Appl Bacteriol 30: 444–449.

Wehrli, E., Scherrer, P., and Kübler, O. (1980) The crystalline layers in spores of *Bacillus cereus* and *Bacillus thuringiensis* studied by freeze-etching and high resolution electron microscopy. European journal of cell biology 20: 283–289.

Wick, R.R., Judd, L.M., Gorrie, C.L., and Holt, K.E. (2017) Unicycler: Resolving bacterial genome assemblies from short and long sequencing reads. PLoS Comput Biol 13: e1005595

Zheng, L.B., Donovan, W.P., Fitz-James, P.C., and Losick, R. (1988) Gene encoding a morphogenic protein required in the assembly of the outer coat of the *Bacillus subtilis* endospore. Genes & development 2: 1047–1054.

Zheng, S., Wolff, G., Greenan, G., Chen, Z., Faas, F.G.A., Bárcena, M., et al. (2022) AreTomo: An integrated software package for automated marker-free, motion-corrected cryo-electron tomographic alignment and reconstruction. J Struct Biol: X 6: 100068.

Zheng, S.Q., Palovcak, E., Armache, J.-P., Verba, K.A., Cheng, Y., and Agard, D.A. (2017) MotionCor2: anisotropic correction of beam-induced motion for improved cryo-electron microscopy. Nat Methods 14: 331–332.

